# Microglia control cerebral blood flow and neurovascular coupling via P2Y12R-mediated actions

**DOI:** 10.1101/2021.02.04.429741

**Authors:** Eszter Császár, Nikolett Lénárt, Csaba Cserép, Zsuzsanna Környei, Rebeka Fekete, Balázs Pósfai, Diána Balázsfi, Balázs Hangya, Anett D. Schwarcz, Dávid Szöllősi, Krisztián Szigeti, Domokos Máthé, Brian L. West, Katalin Sviatkó, Ana Rita Brás, Jean-Charles Mariani, Andrea Kliewer, Zsolt Lenkei, László Hricisák, Zoltán Benyó, Mária Baranyi, Beáta Sperlágh, Ákos Menyhárt, Eszter Farkas, Ádám Dénes

## Abstract

Microglia, the main immunocompetent cells of the brain regulate neuronal function in health and disease, but their contribution to cerebral blood flow (CBF) remained elusive. Here we identify microglia as important modulators of CBF both under physiological conditions and during hypoperfusion. We show that microglia establish direct purinergic contacts with cells in the neurovascular unit that shape cerebral perfusion in both mice and humans. Surprisingly, the absence of microglia or blockade of microglial P2Y12 receptor (P2Y12R) substantially impairs neurovascular coupling in the barrel cortex after whisker stimulation. We also reveal that hypercapnia, which is associated with acidification, induces microglial adenosine production, while depletion of microglia reduces brain pH and impairs hypercapnia-induced vasodilation. Furthermore, the absence or dysfunction of microglia markedly impairs adaptation to hypoperfusion via P2Y12R after transient unilateral common carotid artery occlusion, which is also influenced by CX3CR1-mediated actions. Thus, our data reveal a previously unrecognized role for microglia in CBF regulation with broad implications for common neurological diseases.

## Introduction

Microglia are key regulators of inflammatory processes in the brain and altered microglial activity is linked to the development of common brain diseases such as epilepsy, stroke, Alzheimer’s disease or age-related cognitive decline ^1, 2^. In recent years, the contribution of microglial actions to diverse physiological processes, including brain development, synaptic plasticity, learning or memory has become increasingly recognised ^3, 4^. The complexity of these processes requires dynamic and tightly regulated interactions between microglia and other cell types in the brain, which are incompletely understood. Among these, microglia-neuron interactions are the best documented, while the importance of crosstalk between microglia and astrocytes has recently been established ^5, 6^. Communication between microglia and other cell types is mediated by motile microglial processes, through which microglia perform dynamic surveillance of their environment ^7–9^. Microglial processes are recruited to blood vessels and to other structures in the neurovascular unit (NVU), although the function of these interactions remained vaguely defined ^10, 11^. Microglia-vascular interactions are present in the brain from early development into adulthood, through which microglia regulate blood brain barrier (BBB) permeability, leukocyte extravasation and angiogenesis^10, 12, 13^. In fact, microglia originate from common erythro-myeloid progenitors in the yolk sac ^14^ and are associated with developing blood vessels in the neuroepithelium, while *Pu.1^−/−^* mice or *Csf1^op/op^* mice that lack microglia and macrophages display impaired angiogenesis in the retina ^10, 15^. In the adult brain, CD206-positive perivascular macrophages (PVMs) remain closely associated with blood vessels, whilst CD206-negative microglial cell bodies occupy the brain parenchyma isolated by the glia limitans ^16–18^. PVMs have recently been shown to play an important role in neurovascular dysfunction associated with hypertension, via promoting BBB permeability ^19^. However, little is known about the possible contribution of microglia to vascular dysfunction or perfusion deficits in the adult brain.

Microglia respond to vascular injury, as seen in experimental models of stroke, Alzheimer’s disease or multiple sclerosis ^10, 11^, and microglial processes are recruited to sites of BBB leakage within minutes ^12, 13, 20^. However, we could not detect major differences in the extent of BBB injury after experimental stroke in the absence of microglia ^21^. Interestingly, changes in microglial process dynamics around capillaries are proportional to the level of cerebral blood flow (CBF) reduction during transient ischemia ^22^, suggesting a possible role for microglia-vascular interactions beyond vascular injury.

We have recently identified specific sites on neuronal cell bodies through which microglia shape neuronal responses in health and disease via purinergic mechanisms ^23^. Since microglia interact with both neurons and blood vessels ^21^ and neuronal activity-dependent changes in CBF are precisely controlled via multi-cellular interactions in the NVU ^24^, we argued that microglia are ideally positioned to sense and influence neurovascular responses under normal conditions or when blood flow is disturbed. Thus, we hypothesized that microglia-specific interventions may alter CBF responses to physiological neuronal activation or during hypoperfusion, which – in parallel with inflammatory changes and altered microglial activity – often precedes symptom onset in common neurological disorders ^24, 25^. Here, by using microglia depletion, transgenic mice and pharmacological interventions, we identify microglia as important regulators of CBF during neurovascular coupling, hypercapnia-induced vasodilation and adaptation to hypoperfusion in the cerebral cortex.

## Results

### Microglia form dynamic purinergic contacts with endothelial cells and other cell types in the NVU that regulate CBF

We first investigated the formation and dynamics of microglia-vascular interactions using *in vivo* two-photon imaging. Intravenous FITC-Dextran administration in CX3CR1^tdTomato^ microglia reporter mice allowed 3D reconstruction of penetrating arterioles in the cerebral cortex down to 600 μm below the dura mater (Fig.1a). *In vivo* imaging revealed microglia ensheathing arterial bifurcations at the level of 1^st^, 2^nd^ and 3^rd^ order vessels and identified contacting microglial processes at all levels of the vascular tree (Fig.1a,b and Supplementary Video 1). The average lifetime of contacts ranged between 5 to 15 min and microglial processes frequently re-contacted the same sites at both arterioles and microvessels, suggesting that specific sites for microglia-vascular interactions may exist in the brain. Next, we studied the formation of physical contact between microglia and other cells in the NVU, using the microglial marker P2Y12 receptor (P2Y12R), which is not expressed by any other cells in the brain^26^. Surprisingly, we found that processes of parenchymal microglial cells were extended beyond GFAP-positive perivascular glial endfeet at the level of penetrating arterioles, directly contacting SMA-positive smooth muscle cells (Fig.1c) and endothelial cells in both large vessels and microvessels as evidenced by both confocal- and immunoelectron microscopy (Fig.1d-e). 3D analysis of Z-stacks recorded by confocal laser scanning microscopy (CLSM) revealed that 85% of randomly selected blood vessel segments are contacted by microglial processes and 15% of the endothelial cell surface is covered by microglial processes in the cerebral cortex of mice (Fig.1f). The vast majority of pericytes (83%) also received direct microglial contact (Fig.1g-h). Since ATP (and ADP) is a major chemotactic factor for microglia via P2Y12R ^9^ and purinergic signaling markedly influences CBF in endothelial cells and pericytes ^27^, we turned to electron tomography to investigate the relationship between microglial P2Y12R localization and the vasculature at the highest possible resolution, in 3D. We found contacting P2Y12R-positive microglial processes in close apposition with endothelial mitochondria, while immunogold particles were enriched at the interface (Fig.1i). Unbiased immunofluorescent analysis revealed 214% higher TOM20 immunofluorescence (a mitochondrial marker) in endothelial cells at microglial contact sites (Fig.1j and Supplementary Fig.1d). These results suggest that similarly to that seen at somatic purinergic junctions ^23^, ATP released at the perivascular compartment could also act as a chemotactic signal for microglial processes. Importantly, immunoelectron microscopy also confirmed the direct contact between P2Y12R-positive microglial processes and endothelial cells in the human brain (Fig.1k). The perivascular endfeet of GFAP expressing astrocytes contribute to CBF regulation ^28^. We found that 93% of astrocytes were contacted by P2Y12R-positive microglial processes, while microglial cell bodies were found directly attached to 18% of astrocytes (Fig.1l and Supplementary Fig.1a-b). To visualize perivascular astrocyte endfeet, Aquaporin-4 (AQP4) immunostaining was used, showing that microglial processes directly contact endothelial cells at sites void of the AQP4 signal (Supplementary Fig.1c) or by extending through the astrocytic endfeet layer (Supplementary Fig.1e). Combined immunogold-immunoperoxidase labeling and electron microscopy confirmed the direct contact between microglial processes and parenchymal astrocytes or perivascular astrocytic endfeet (Fig.1m). Similar observations were made in the human cerebral cortex from both aged and middle-aged patients who died in non-neurological conditions: P2Y12R-positive microglial processes established contact with both perivascular astrocyte endfeet and the endothelial monolayer of small arterioles and capillaries (Fig.1n-o and Supplementary Fig.1f). Furthermore, we found that individual microglial cells contact multiple microvessels and nearby neurons simultaneously in the brain (Fig. 1p). Thus, microglial processes not only directly contact cells in the NVU along the vascular tree (Supplementary Fig.1g), which are known to shape CBF ^24, 25, 29–31^, but simultaneous contacts with neurons and vascular structures may provide an ideal opportunity for microglia to influence neurovascular responses.

**Figure 1.**
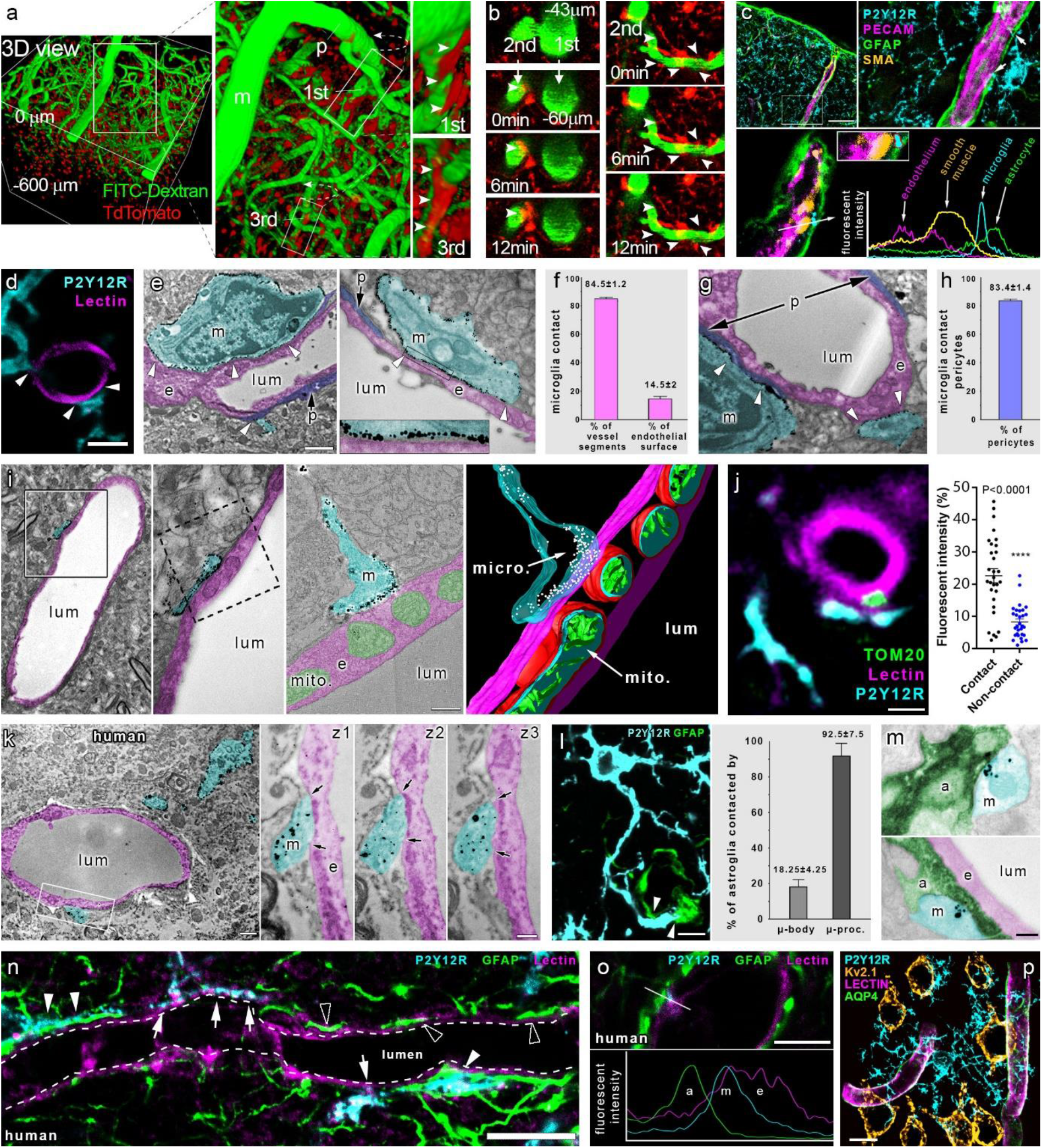
Microglia form physical contact with cells in the neurovascular unit that regulate cerebral blood flow. **a)** 3D reconstruction of *in vivo* two-photon Z-stacks down to 600 μm below the dura mater in the cerebral cortex of CX3CR1^tdTomato^ mice. Meningeal (m), penetrating (p), and 1^st^ to 3^rd^ order capillaries have been identified for *in vivo* time-lapse imaging. Note contacting microglia (arrowheads) at different vascular beds. **b)** Real-time imaging reveals microglial processes (arrowheads) contacting different segments of the vascular tree (visualized by i.v. FITC-Dextran). **c)** Microglial processes are extended beyond the perivascular glial endfeet and form direct contact with smooth muscle cells (arrows) at the level of penetrating arteries. **d)** CLSM images show microglia (P2Y12R, cyan) contacting endothelial cells (Tomato lectin, magenta) in the cerebral cortex. **e)** EM images show microglia (m, P2Y12R-immunogold labeling, cyan) directly contacting endothelial cells (e, magenta) and pericytes (p, purple). **f)** Frequency of vessels receiving microglial contact, and microglial process coverage of endothelial cell surface. **g)** EM images show microglia (m, P2Y12R-immunogold labeling, cyan) directly contacting pericytes (p, purple). **h)** 83.4 ± 1.4% of pericytic cell bodies are contacted by microglial processes. **i)**3D reconstruction of electron tomogram shows clustering of anti-P2Y12R-immunogold on microglial processes (m) directly contacting the endothelium (e) of an arteriole / post-capillary venule. Left two panels are conventional EM images of the same area on the adjacent ultrathin section. The right panels show tomographic virtual section and 3D reconstruction of the direct contact. **j)** Unbiased anatomical analysis reveals an enrichment of endothelial mitochondria (TOM20+, green), at sites of microglial contacts (P2Y12R+, cyan). For details of analysis refer to Supplementary Fig. 1D and the Methods. **k)** EM images show microglia (m, P2Y12R-immunogold, cyan) directly contacting endothelial cells (e, magenta) in human neocortex. z1-z3 panels show the contact on three consecutive ultrathin sections, arrows mark the edges of direct membrane contact. **l)** CLSM image show microglia (P2Y12R, cyan) contacting the cell body of an astrocyte (GFAP labeling, green) and astrocytic endfeet (arrowheads). 92.5±7.5% of astrocytes are contacted by microglial processes. **m)** EM images show direct contact between microglial (m, cyan) and astrocytic (a, green) processes. (e-endothelial cell, magenta; lum-lumen). **n)** CLSM images in human neocortex reveal P2Y12R+ microglial processes (cyan) contacting perivascular astrocytes (GFAP, green) on astrocyte endfeet (white arrowheads) and endothelial cells (tomato-lectin, magenta, white arrows), with astrocytic endfeet directly touching the endothelial monolayer (empty arrowheads). **o)** CLSM image and fluorescent intensity plots show microglial process (m) contacting the endothelial layer (e) within the astrocytic layer (a). **p)** CLSM maximal intensity plot shows individual microglial cells (cyan) contacting several microvessels (lectin-magenta, GFAP-green) and neurons (ocker) simultaneously. Scale bars: **c**, 50 μm; **d,**3 μm; **e-g**, 2 μm on e-left panel, 500 nm for e and g; **i**, 200 nm; **j**, 2 μm; **k**, 1 μm on the left panel and 400 nm on z3; **l,** 10 μm; **m,** 200 nm; **n,** 10 μm; **o,** 5 μm; **p,** 10 μm.

### Microglia contribute to neurovascular coupling via P2Y12R-mediated actions

In our previous studies, we could not detect major alterations in the number or morphology of endothelial cells, astrocytes or pericytes after elimination of microglia by PLX5622 ^21^. However, we found it important to investigate whether prolonged absence of microglia could compromise overall brain perfusion or metabolism. To this end, we performed HMPAO-SPECT and FDG-PET measurements ^32, 33^ after microglia depletion by PLX5622 ^34^. No significant changes in HMPAO or FDG uptake were observed in any brain areas after microglia depletion (Supplementary Fig. 2a-c). Next, we aimed to investigate the role of microglia in CBF responses to physiological stimuli. In the adult brain, CBF not only varies in proportion to the energy consumption, but increased neuronal activity leads to increases in CBF, highly restricted to the activated areas, a phenomenon called functional hyperemia ^2^. Thus, we turned to the whisker-stimulation model, which is widely used to study the mechanisms of neurovascular coupling in mice ^35^ and performed laser speckle contrast imaging (LSCI), a highly sensitive approach optimized to assess changes in the microcirculation through the intact skull bone in real time ^36^. Whiskers on the left side were stimulated under mild ketamine-medetomidine sedation, allowing stable and reproducible CBF responses to be observed (Fig.2a-b). Surprisingly, the absence of microglia resulted in impaired functional hyperemia, as evidenced by a significant, 15.3% smaller CBF response in the right barrel cortex compared to that seen in control mice after a series of stimulation (6 series of stimulations for 30s each; Fig.2b-c and Supplementary Video 2). Supporting the role of microglia in neurovascular coupling, a 17% smaller CBF response to whisker stimulation was seen after acute microglial P2Y12R blockade by a specific P2Y12R inhibitor, PSB-0739, injected into the cisterna magna 35 min prior to LSCI measurements^23^. Because only microglia express P2Y12R in the brain ^23, 37^, this way we could also validate the specificity of microglial actions on CBF responses. To extend these observations with genetic P2Y12R blockade, another series of measurements were performed, using manual whisker stimulation followed by a set of electromechanically controlled stimulations. We found reduced CBF responses to whisker stimulation in both microglia-depleted and P2Y12R KO mice, irrespective of the type of stimulation used (Fig. 2d-f). To also investigate the effect of microglia depletion on cortical CBF responses to whisker stimulation by an alternative approach, we turned to functional ultrasound (fUS) imaging measurements, which detect hemodynamic changes in the brain based on cerebral blood volume (CBV) ^38^. We found that the absence of microglia resulted in significantly smaller (by 28%) CBV increases in the contralateral barrel cortex in response to whisker stimulation compared to that seen in control mice (Fig.2g-i). Thus, microglia- and microglial P2Y12R-mediated actions are important to maintain normal blood flow responses to neuronal activity induced by physiological stimuli in the cortical microcirculation.

**Figure 2.**
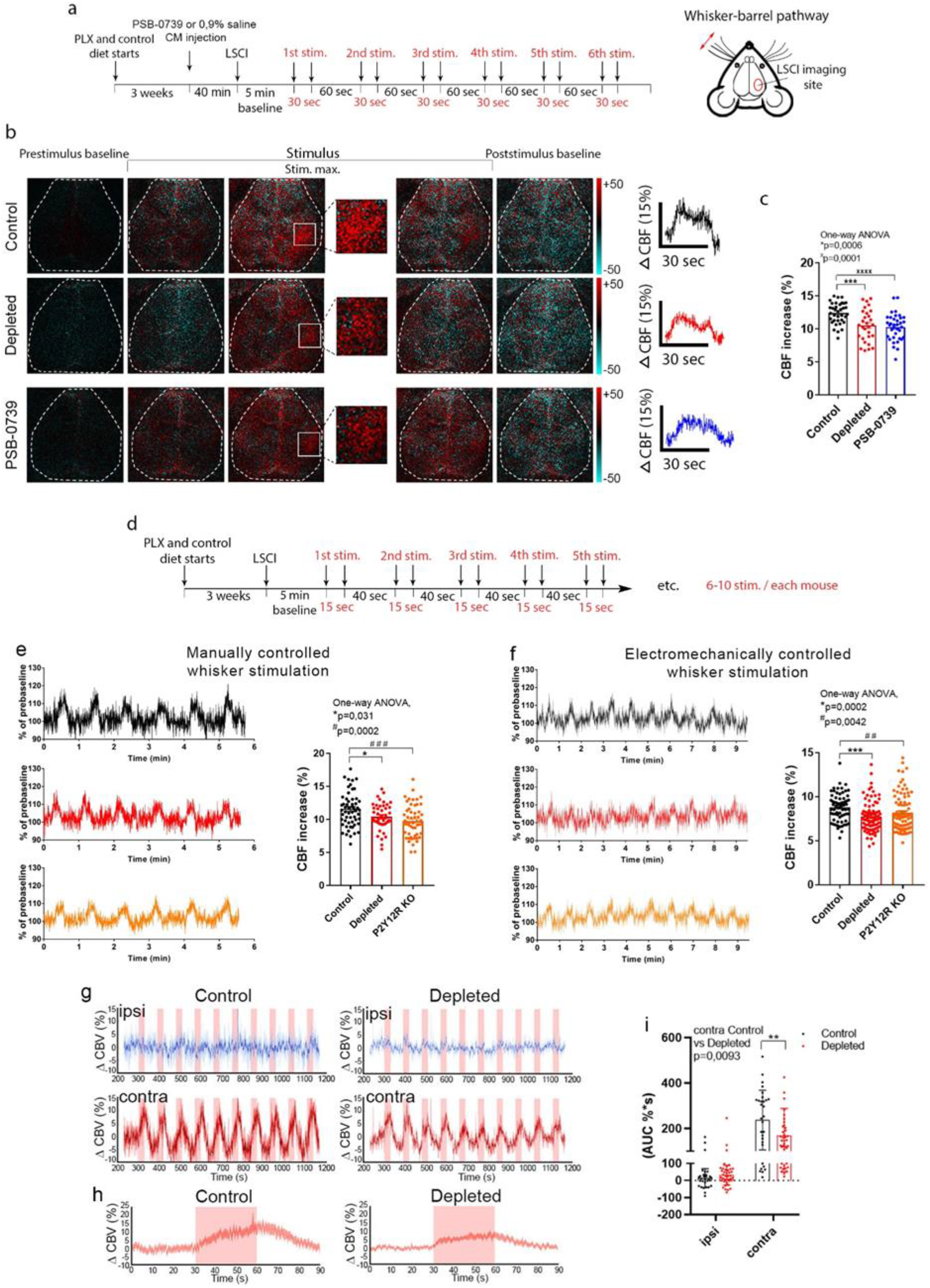
Microglia contribute to neurovascular coupling in P2Y12 receptor-mediated manner. **a)** Schematic showing the outline of the experiment. Microglia were depleted by PLX5622. Mice were injected either with saline (Control) or with PSB-0739 into the cisterna magna 40 min prior to LSCI imaging. CBF changes to whisker stimulation were examined in the contralateral barrel cortex. **b)** Difference images show CBF changes in the right barrel cortex relative to baseline in response to contralateral whisker stimulation before, during and after stimulus (white rectangle indicates the barrel field). Time course of stimulus-evoked CBF responses is shown in the right of panel b. **c)** Absence of microglia or acute blockade of P2Y12R reduce the maximum of evoked CBF responses compared to the control. **d)** Protocol of manually and electromechanically controlled whisker stimulation. **e,f)** Representative CBF traces and quantification show impaired neurovascular coupling response in the absence of microglia and in P2Y12R KO mice compared to control. **g**) Functional ultrasound (fUS) imaging also show reduced CBF (CBV) responses as compared to controls in the ipsilateral (ipsi) and contralateral (contra) barrel cortex. Representative traces of 10 subsequent stimulations (30s each) are shown for control and microglia-depleted mice. **h**) Peak trace averages of the contralateral side in control and depleted mice, with 95% confidence intervals. **i**) Averaged area under the curve (AUC) distribution for each group, as shown in pink window on panel h). Data are expressed as mean ± SEM; n=36, n=29, n=36 stimulations from 6 control, 5 depleted and 6 PSB-0739 mice, respectively (c); n=57, n=45, n=54 stimulations from 7 control, 6 depleted and 7 P2Y12R KO mice, respectively (e), n=69, n=81, n=92 stimulations from 7 control, 9 depleted and 9 P2Y12R KO mice, respectively (f); n=30 and n=40 stimulations from 3 control and 4 depleted mice, respectively (i). *p control vs depleted, ^×^p control vs PSB-0739 injected mice, #p control vs P2Y12R KO mice, one-way ANOVA followed by Dunnett’s multiple comparison test (c,e), Kruskal-Wallis test followed by Dunn’s multiple comparison test (f), two-way ANOVA followed by Sidak’s multiple comparisons test (i).

### Changes in whisker stimulation-evoked neuronal responses do not explain altered CBF responses after microglia manipulation

To test whether substantial shifts in neuronal activity could explain marked CBF differences seen upon microglia manipulation, we performed unilateral whisker stimulation under identical ketamine-medetomidine anesthesia as above, while recording neuronal activity from the contralateral barrel cortex, either using chronically implanted tetrode electrodes or *in vivo* two-photon calcium imaging. We isolated n=42, n=41 and n=61 putative single units from 2 electrophysiological recordings of each control, microglia-depleted and P2Y12R KO mice respectively (n = 5). This allowed us to test baseline firing rates in the stimulus-free periods as well as stimulus-induced firing responses of individual neurons. We found significantly increased baseline firing rates of barrel cortex neurons in both microglia-depleted and P2Y12R KO mice compared to controls (Fig. 3a-d). However, either using electromechanically controlled automated whisker stimulation or manual stimulation, we did not detect differences in the extent of stimulus-evoked neuronal responses (Fig. 3e). Then, we turned to *in vivo* two-photon measurements, using electromechanical whisker stimulation, which was repeated two times with 40s intervals (Fig. 3f). Only neurons in the contralateral barrel cortex, which specifically responded to both stimuli were selected for analysis. We found that somatosensory stimulus-induced increases in the neuronal GCaMP6s signal in Thy1-GCaMP6s mice did not reveal significant differences between control and microglia-depleted mice (Fig.3g and Supplementary Video 3; dF/F AUC 1517±677 vs 1481±987 in control and depleted mice, respectively). Thus, while the absence (PLX5622-depleted) or dysfunction (P2Y12R KO) of microglia may shift baseline neuronal activity, the above measurements collectively suggest that alterations in stimulus-evoked neuronal responses do not explain the marked differences in CBF changes observed after microglia manipulation.

**Figure 3.**
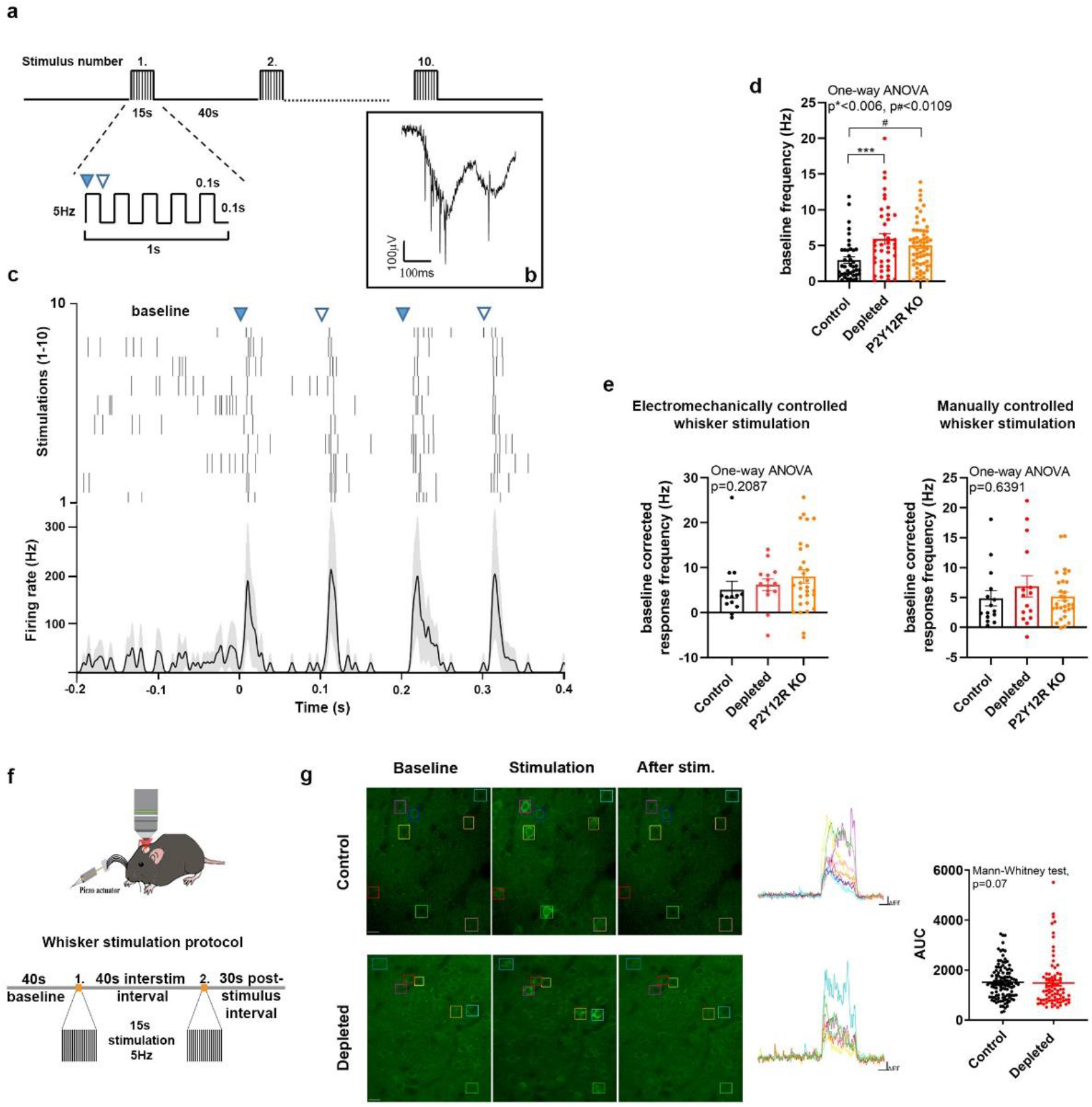
Whisker stimulation-evoked neuronal responses in the barrel cortex do not explain altered CBF responses after microglia manipulation. **a)** Schematics of the whisker stimulation protocol. Whiskers were stimulated electromechanically with 5 Hz, causing alternating passive deflections of the vibrissae in the anterior and posterior directions (filled and empty arrowheads, respectively) for 15s, followed by a 40s pause, repeated 10 times. **b)** Raw tetrode data showing extracellular spikes recorded from the barrel cortex. **c)** Example of a single neuron activated by passive whisker deflections. Top, raster plot aligned to whisker stimulation onset (black ticks, individual action potentials). Bottom, peri-stimulus time histogram showing mean firing responses of the same neuron (shading, SEM). **d)** Baseline firing rates were significantly higher in depleted and P2Y12R KO mice compared to controls. Data are presented as mean ± SEM, n=4 control, n=3 depleted and n=5 P2Y12R KO mice, p*** control vs depleted, p# control vs P2Y12R KO, one-way ANOVA with Dunett’s multiple comparisons test. **e)** Stimulus-induced firing rate changes were comparable across controls and microglia depleted mice using either electromechanically or manually controlled whisker stimulation. Data are presented as baseline corrected response frequency (for corresponding baseline frequencies, mean ± SEM), n=4 control, n=3 depleted and n=5 P2Y12R KO mice, Kruskal-Wallis test with Dunn’s multiple comparisons test. **f)** Schematic outlines of the whisker stimulation protocol used for *in vivo* two-photon [Ca^2+^]_i_ imaging in the barrel cortex of Thy1-GCaMP6s mice. **g)** Representative images show stimulus-evoked neuronal [Ca^2+^]_I_ responses with individual traces of neurons labelled with rectangles during baseline imaging, 15s stimulation and after stimulation. AUC (area under the curve) values of neuronal GCaMP6s signal changes in response to electromechanically controlled whisker stimulation (p=0.07, Mann-Whitney test) in control and microglia-depleted mice. Data are presented as mean ± SEM, n=56 neurons from control and n=40 neurons from depleted mice from two trials, Mann-Whitney test, n=4 mice per group.

### Microglia shape hypercapnia-induced vasodilation in a P2Y12R-dependent manner

While our neurovascular coupling studies above identified microglia as an important cell type to shape CBF via P2Y12R, we aimed to further investigate the mechanisms involved, using hypercapnic challenge to induce vasodilation independently of direct neuronal stimulation. Hypercapnia is considered to induce primarily endothelium-driven vasodilation and related increase in CBF, although actions of astrocytes and other cells also appear to be important for this process ^28, 39–42^. To study whether microglia respond to hypercapnic challenge, we first performed *in vivo* two-photon imaging in CX3CR1^tdTomato^ mice after intravenous FITC-Dextran administration. We found that a population of arteriole-associated, dynamic microglial processes readily changed their morphology in response to vasodilation both at arterioles and microvessels due to 2 min inhalation of 10% CO_2_ under normoxic conditions (Fig.4a and Supplementary Fig.3a). Around arterioles, SR101-labeled perivascular astrocyte endfeet were dynamically contacted by CX3CR1^GFP/+^ microglia (Fig.4b) and the number of contacting phylopodia at the end of microglial processes increased in response to hypercapnia (Fig.4b). Importantly, *in vivo* two-photon imaging revealed significantly impaired hypercapnia-induced vasodilation in meningeal and penetrating arteries (Fig.4c-d), which paralleled smaller CBF responses in microglia-depleted mice as assessed by LSCI (Fig.4c,e-g). To exclude the potential effect of alpha2 adrenergic blockade via the cardiovascular system during ketamine-medetomidine anesthesia^43^, we repeated hypercapnic challenge after the administration of atipamezole, an alpha2 receptor antagonist^44^. Hypercapnia-induced CBF response was similarly reduced in microglia-depleted mice compared to controls in the presence of atipamezole (Fig.4h). Importantly, baseline and hypercapnia-induced pCO_2_, pO_2_ levels and pH in blood samples taken from the femoral artery were not altered by microglia depletion (Supplementary Fig.3b). We also investigated whether elimination of P2Y12R would alter hypercapnia-induced vasodilation. *In vivo* two-photon imaging revealed a 37% smaller vasodilation in response to hypercapnia in P2Y12R-deficient double transgenic (CX3CR1^GFP/+^ × P2Y12R KO) mice as compared to control (CX3CR1^GFP/+^) mice (Fig.4i and Supplementary Video 4). In line with these observations, neuronal activity did not differ between control, microglia-depleted and P2Y12R KO mice during hypercapnic challenge (Fig. 4j).

**Figure 4.**
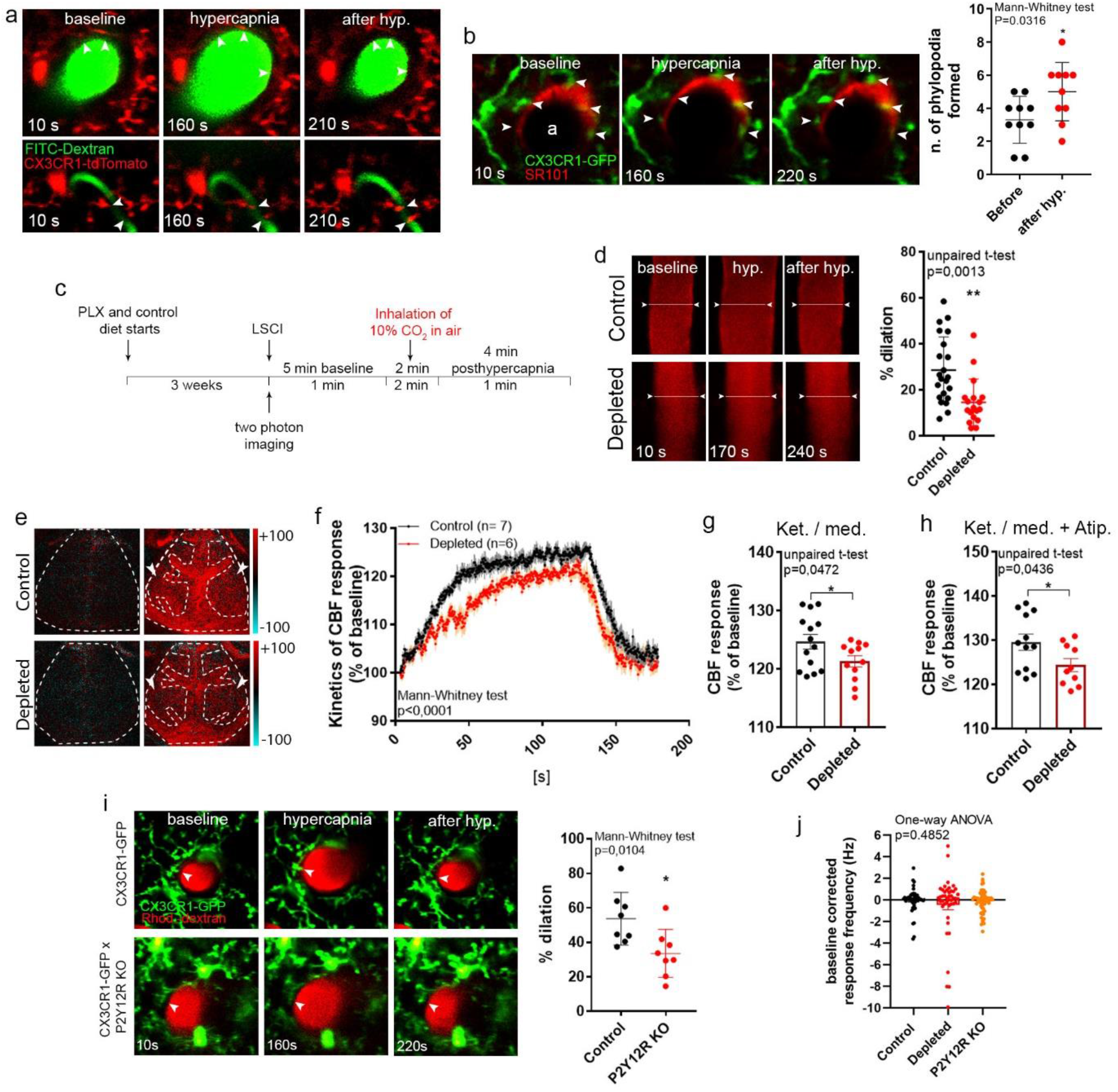
Microglia contribute to hypercapnia-induced vasodilation. **a)** *In vivo* two-photon resonant (32Hz) imaging was performed in the somatosensory cortex of CX3CR1^tdTomato^ mice during hypercapnia (by inhalation of 10% CO_2_ under normoxic conditions). The middle panel shows the maximal vasodilation provoked by hypercapnia. **b)** Identical hypercapnic challenge and imaging protocol was performed in CX3CR1^GFP/+^ mice after intracortical injection of SR101 to visualize astrocytes. The number of phylopodia formed at the end of contacting microglial processes (arrowheads) increased in response to hypercapnia. **c)** Experimental protocol of hypercapnic challenge for *in vivo* two-photon imaging (a-b,d,i) and LSCI (e-h). **d)***In vivo* two-photon imaging reveals impaired vasodilation at the level of penetrating arteries in the absence of microglia. **e)** Difference images show reduced CBF response in microglia depleted mice to hypercapnic challenge (ROIs are labelled with arrowheads). **f)** The average kinetics of hypercapnic responses show difference in depleted mice compared to controls. **g)** Hypercapnia-evoked CBF response is markedly decreased in the absence of microglia under ketamine-medetomidine or **h)** ketamine-medetomidine anesthesia after administration of atipamezole. **i)** Elimination of P2Y12R impairs hypercapnia-induced vasodilation in double transgenic (CX3CR1^GFP/+^ × P2Y12R KO) mice compared to P2Y12R-competent CX3CR1^GFP/+^ mice. **j)** During hypercapnic challenge neuronal activity did not differ between control, microglia-depleted and P2Y12R KO mice. Data are expressed as mean ± SEM. n=5 control and n=5 depleted mice (b) n=22 and n=18 vessels from 8 control and 6 depleted mice, respectively (d). n=14-12 ROIs from 7 control and 6 depleted mice, 2 ROIs/mouse (f,g); n=12-11 ROIs from 6 control and 5 depleted mice, 2 ROIs/mouse (h); n=8-8 vessels from n=5 control and n=5 P2Y12R KO mice (i); n=49 single units/cells in control, n=44 in depleted and n=61 in P2Y12R KO group (j). Mann-Whitney test (b,f,i), unpaired t-test (d,g,h) or Kruskal-Wallis test with Dunn’s multiple comparison (j).

### Selective elimination of microglia results in impaired adaptation to cerebral hypoperfusion

Next, we aimed to investigate the impact of microglial responses on CBF during hypoperfusion, which has high pathophysiological relevance for diverse vascular diseases including stroke, chronic hypoperfusion or vascular dementia among others ^24, 25, 45^. Because microglial activity is altered in response to hypoxia or cortical hypoperfusion ^22, 46^, we developed a model to study the actions of hypoperfusion-primed microglia on subsequent CBF changes by repeating transient unilateral common carotid artery occlusion (CCAo) and reperfusion 3 times (Supplementary Fig.4a-b). Redistribution of blood flow to the ipsilateral cortical circulation requires vasodilation ^47^, and unilateral CCAo does not cause cerebral ischemia ^47, 48^, therefore this model was found also useful to study vascular adaptation responses in the absence of neuronal injury, which is influenced by microglia manipulation ^21^. We found that blood vessel-associated microglial processes respond to CBF drop, as shown by increased process motility immediately after CCAo during *in vivo* two-photon imaging (Fig.5a). These changes were similar in CX3CR1^tdTomato^ and CX3CR1^GFP/+^ mice and high-resolution automated analysis demonstrated that alterations in microglial process morphology are maintained up to 24h after CCAo (Fig.5b). Importantly, LSCI measurements showed markedly impaired adaptation to cortical hypoperfusion in the absence of microglia. This was evidenced by lower baseline-corrected CBF values after 5min CCAo and subsequent reperfusion for 5min, which effect gradually increased as CCAo and reperfusion were repeated two more times (p<0.0001, two-way ANOVA, Fig.5c-d and Supplementary Fig.4a-b). In fact, average CBF values by the third occlusion reached only 79% of baseline in microglia depleted mice, as opposed to 89% in control mice in the ipsilateral hemisphere (Fig.5d; and Supplementary Video 5). Interestingly, the absence of microglia also markedly impaired CBF recovery after repeated CCAo in the contralateral hemisphere (p<0.0001, two-way ANOVA, Fig.5d; and Supplementary Video 5), indicating that microglial actions are involved in normalizing CBF responses during hypoperfusion. Impaired CBF recovery was also evident in both hemispheres between the 2^nd^ and the 3^rd^ reperfusions in microglia-depleted mice (2.8 fold larger reduction compared to control mice both ipsilaterally and contralaterally, p=0.042 and p=0.048 respectively, unpaired t-test).

**Figure 5.**
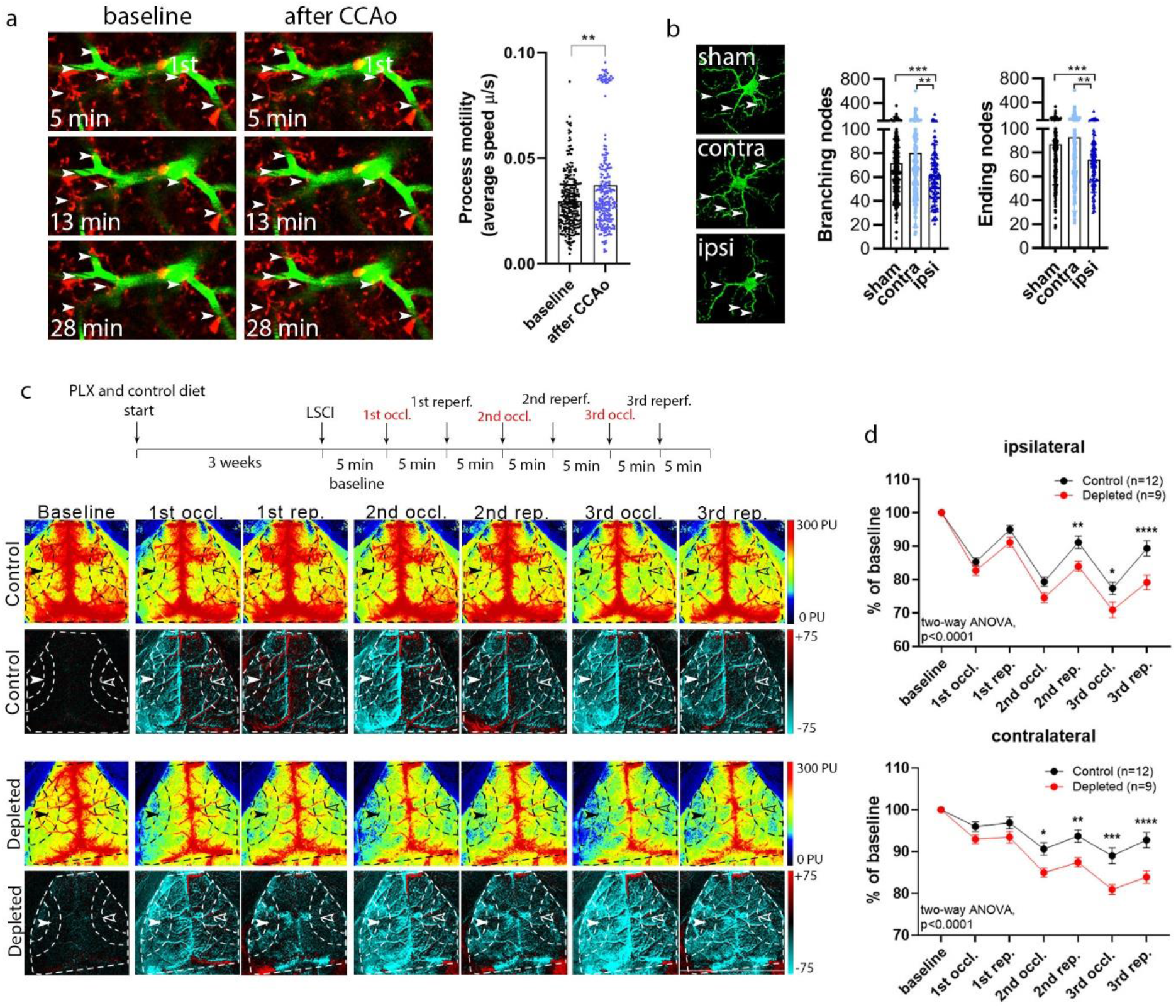
Adaptation to cortical hypoperfusion is impaired in the absence of microglia. **a)** *In vivo* two-photon imaging reveals increased microglial process motility (arrowheads) to repeated (3x) CCAo in CX3CR1^tdTomato^ mice (1^st^ denotes first-order capillary). **b)** Automated morphological analysis demonstrates reduced number of branching-, and ending nodes of microglial processes ipsilaterally in CX3CR1 ^GFP/+^ mice 24h after 3x CCAo compared to the contralateral side (contra) and sham animals in the cerebral cortex. **c)** Schematics of the experimental protocol of 3x CCAo. Microglia were selectively eliminated from the brain by PLX5622. Repeated CCA occlusions were performed in control and microglia depleted mice by pulling away the CCA with a silk thread. Representative perfusion (1^st^ and 3^rd^ rows)-, and difference LSCI images (2^nd^ and 4^th^ rows) showing cortical perfusion changes in response to 3x CCAo in control and microglia depleted mice. Dashed lines indicate the MCA2 territory in both the ipsilateral (white arrowheads) and contralateral (empty arrowheads) hemisphere corresponding to the quantification shown in panel d. Venous sinuses were excluded from the analysis. **d)** CBF responses of the MCA2 area (as shown by arrowheads) to 3x CCA occlusion. CBF responses are shown as percentage of baseline. A significant CBF reduction is seen in the absence of microglia in both hemispheres. Data are expressed as mean ± SEM. n=9-12 mice, *p control vs depleted, Mann-Whitney test (a); branching-ending nodes of n=386-388 sham, n=197 contralateral n=134 ipsilateral cells from n=3 sham and n=3 CCAo mice. Kruskal-Wallis test followed by Dunn’s multiple comparisons test (b); two-way ANOVA followed by Sidak’s multiple comparison test (d),.

### Microglial actions on CBF are partially CX3CR1-dependent and differ from those of perivascular macrophages

Next, we performed the repeated CCAo paradigm in CX3CR1^GFP/GFP^ (fractalkine receptor-deficient) mice, which show changes in a broad spectrum of microglial properties and are widely used to study microglia-dependent mechanisms (Fig.6) ^49^. Interestingly, heterozygous CX3CR1^GFP/+^ mice, which have reduced fractalkine receptor levels showed impaired blood flow recovery after repeated CCAo, while adaptation to hypoperfusion was improved in CX3CR1^GFP/GFP^ mice (Fig.6a-c). These results suggested that compensatory actions by another cell type may interfere with the effects of microglial CX3CR1 on blood flow regulation. In fact, when studying the anatomical relationship between CX3CR1-positive microglia and brain microvessels, we noticed that a subpopulation of CD206-positive PVMs also express the fractalkine receptor (in the cerebral cortex, 2.5% of CD206+ cells were strongly, 20% of CD206+ cells were weakly CX3CR1-GFP immunopositive, Fig.6d-f). While P2Y12R^+^ / CX3CR1^+^ / CD206^−^ microglial cell bodies reside in the brain parenchyma, P2Y12R^−^ / CX3CR1^+^ / CD206^+^ PVMs are located near the endothelial monolayer outside the glia limitans (Fig.6 d-f). To investigate whether PVMs could be in part responsible for the CBF changes seen in the present CCAo model, we selectively eliminated these cells by ICV administration of clodronate without an effect on resident microglia (Fig.6g and Supplementary Fig.5). The absence of PVMs did not influence blood flow responses after repeated CCAo (Fig.6h). Thus, microglia-mediated effects on cortical perfusion differ from those induced by PVMs, while complete absence of the fractalkine receptor may lead to compensatory changes in neurons ^50^ or cells other than PVMs.

**Figure 6.**
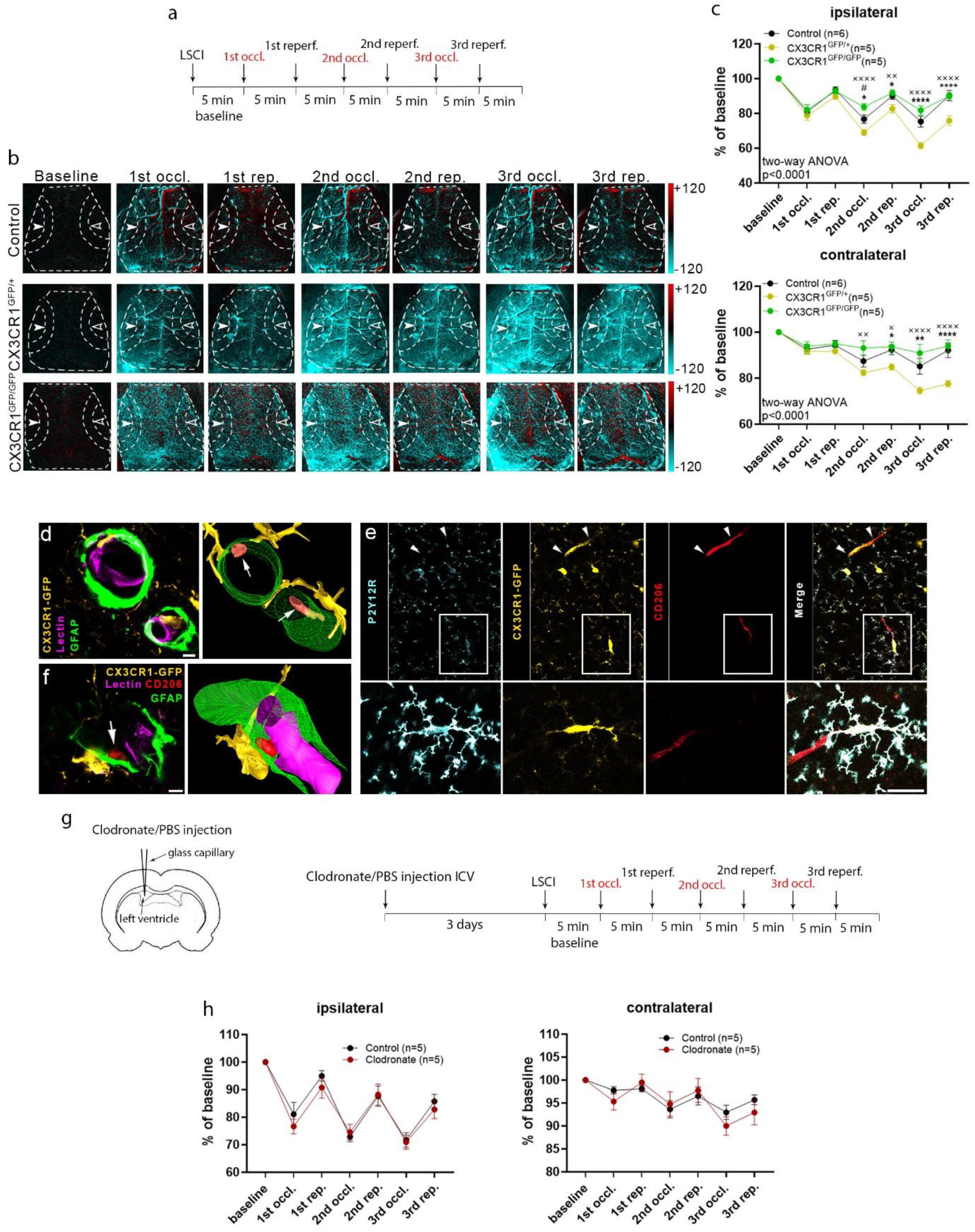
Microglial actions on CBF take place via CX3CR1 signaling and are distinct from the effects mediated by perivascular macrophages. **a)** Outline of the experimental protocol. CCAo was performed with a silk thread. **b)** Representative difference images show altered haemodynamic responses in CX3CR1^GFP/+^ and CX3CR1^GFP/GFP^ mice in both hemispheres. Dashed lines show the MCA2 area both in the ipsilateral (white arrowheads)-, and in the contralateral hemisphere (empty arrowheads) corresponding to quantitative analysis on panel c. **c)** CX3CR1^GFP/+^ mice show significantly impaired CBF recovery in both hemispheres from the 2^nd^ CCA occlusion compared to control and to CX3CR1^GFP/GFP^ mice. However, no significant difference is seen between CX3CR1^GFP/GFP^ and control mice. n=6-5-5 mice. **d)** Cross section CLSM image (left) and 3D reconstruction (right) of two blood vessels, showing CX3CR1^GFP/+^ expressing microglia (yellow) that form contact with tomato lectin-positive endothelial cells (magenta). Microglia and perivascular macrophages (PVMs, between astrocytic endfeet and endothelial layer, orange in 3D model, marked by arrows) are both expressing CX3CR1^GFP/+^ (yellow). **e)** CLSM images show that PVMs (CD206-labeling, red) can be positive for CX3CR1 (colocalisation marked by arrowheads), but negative for P2Y12R (cyan). Inserts show microglial processes contacting a CD206-positive PVM. **f)** CX3CR1^GFP/+^ expressing microglia contact GFAP-labeled astrocytic endfeet. CD206-positive perivascular macrophages reside between astrocytic endfeet and lectin-labeled endothelial cells (magenta). **g)** PVMs were eliminated from the brain by ICV liposomal clodronate injection before LSCI measurements. **h)** No difference in CBF is seen between clodronate-treated and control mice after 3x CCAo. Data are presented as mean ± SEM. n=5-5 mice **\*** p control vs CX3CR1^GFP/+^, ^#^p control vs Cx3CR1^GFP/GFP^, ^×^p CX3CR1^GFP/GFP^ vs CX3CR1^GFP/+^; two-way ANOVA followed by Tukey’s multiple comparison test (c, h). Scale bars: **d**, 5 μm; **e**, 3 μm **f**, 20 μm.

### Microglia respond to hypercapnia by producing adenosine, while depletion of microglia lowers brain pH

To further investigate the mechanisms through which microglia shape CBF, cortical blood flow (by laser Doppler), tissue pH (by using pH-selective electrode) and direct current (DC) potential were simultaneously assessed during hypercapnia and repeated CCAo. Surprisingly, baseline brain pH was significantly lower in depleted mice, while the relative amplitude of the hypercapnia-induced negative pH shift was not different in control animals (Fig. 7a). As seen previously with LSCI (Fig.4e-g), laser Doppler flowmetry showed significantly smaller hypercapnia-induced CBF elevation in the absence of microglia (Fig.7b). Next, we performed 3x CCAo to study how the absence of microglia alters extracellular pH during adaptation to cortical hypoperfusion. For these experiments, CCAo was performed by a mechanical occluder ^51^, to achieve more abrupt, complete cessation of carotid blood flow than in other CCAo studies, where the vessel was pulled away by using a silk thread, resulting in moderate CBF reduction during CCA occlusion (Fig.5d and Fig.6c). Surprisingly, we found that while CBF was reduced by 41% during the first occlusion in control mice, a larger, 81% drop was seen in microglia depleted mice, which triggered spreading depolarization (SD) in 6 out of 6 animals (Fig.7c-d) and progressive oligaemia (Fig.7e), which was not seen in controls. Extracellular pH was also markedly lower in depleted mice during CCAo, with a sharp drop with SD after the first occlusion (Fig.7c,f). Thus, while the absence of microglia promotes acidification in the cerebral cortex, it leads to gradually decreasing CBF during repeated CCAo, which is exacerbated by progressive oligaemia if SD is triggered due to the marked drop of CBF.

**Figure 7.**
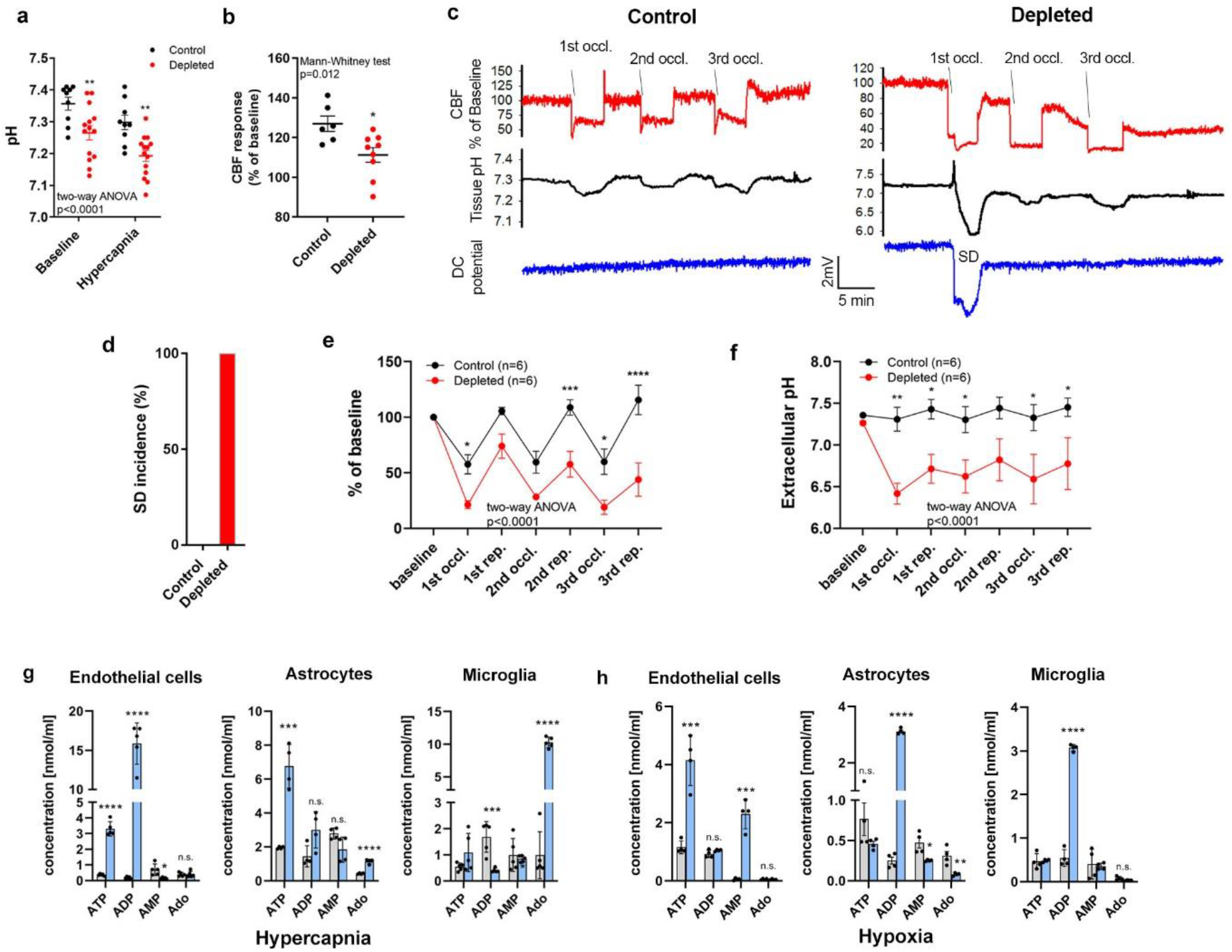
Microglia produce adenosine in response to hypercapnia, while microglia depletion leads to negative pH shift and hypoperfusion. **a)** Depleted mice show reduced extracellular brain pH and **b)** reduced CBF in response to hypercapnia. **c-d:** CBF by laser Doppler flowmetry, tissue pH by pH-selective electrode and DC potential were simultaneously assessed during 3x CCAo performed by a mechanic occluder. The induction of spreading depolarization (SD) after the 1^st^ occlusion was observed in 6 out of 6 microglia depleted mice and none in controls, which was associated with larger negative pH shift and progressive oligaemia by the end of the 3^rd^ reperfusion. **e)** CBF response characteristic to 3x CCAo and **f)** extracellular pH. **g-h)** Effect of hypercapnia (g) and hypoxia (h) on levels of purinergic metabolites in primary endothelial-astrocyte- and microglia cultures as measured by HPLC., Ado, adenosine. Data are expressed as mean ± SEM. n=10 and 16 hypercapnic challenges from 5 control and 8 depleted mice, respectively (a); n=6 control and n=9 depleted mice per group (b); n=6 mice per group (e,f); Mann-Whitney test (b), two-way ANOVA followed by Sidak’s multiple comparison (a,g,e,f,h).

Both hypoxia and hypercapnia may develop during hypoperfusion with hypercapnia driving acidification in the brain ^41^. Because the present studies collectively suggested that microglial actions may counteract cortical hypoperfusion by facilitating CBF recovery through purinergic interactions with both endothelial cells and astrocytes (Fig.1 and Fig.4), we next investigated purinergic metabolites that may mediate these changes. As expected, hypercapnic challenge reduced both extracellular and intracellular pH in astrocyte cultures (Supplementary Fig. 6a-b). HPLC measurements revealed increased ATP and adenosine production by astrocytes, while ATP and ADP were produced by cultured endothelial cells in response to hypercapnia (Fig.7g). Interestingly, ATP, but not ADP was increased in identical endothelial cell cultures after hypoxic challenge, while increased ADP levels were found in astrocyte cultures (Fig.6h, Supplementary Fig.6c). Importantly, hypercapnia, but not hypoxia triggered a robust, ten-fold increase in microglial adenosine production (Fig.7g, h). Collectively, these experiments suggest that hypercapnia and hypoxia lead to rapid production of different purinergic metabolites by endothelial cells and astrocytes, while adenosine is produced by microglia, which also modulate brain pH and contribute to vasodilation in response to different vascular challenges.

### The effect of microglial actions on CBF is mediated via P2Y12R signaling during adaptation to hypoperfusion

Our HPLC studies demonstrated that purinergic mediators are released in a stimulus-specific manner from endothelial cells and astrocytes during hypercapnia and hypoxia, which both occur during hypoperfusion. ADP is the main ligand for microglial P2Y12R, which can rapidly form upon ATP hydrolysis by ectoATPases, expressed by microglial processes ^23^ among other cells. To further explore the mechanisms through which microglia contribute to the regulation of cortical perfusion, we tested whether an inhibition of microglial P2Y12R using either P2Y12R KO mice or acute blockade by PSB-0739 injected into the cisterna magna^23^, alters CBF responses after repeated CCAo. In these experiments, similarly to most studies excluding those presented in Fig.7, CCAo was performed by a silk thread to induce moderate hypoperfusion that did not trigger SD as clearly shown by LSCI measurements. Importantly, we found that blood flow recovery during repeated CCAo was markedly impaired after both genetic and pharmacological P2Y12R blockade in the ipsilateral and in the contralateral hemispheres (Fig.8a-c), similar to that seen in microglia depletion studies (Fig.5). Thus, these results collectively suggest that both microglia and microglial P2Y12R are essential for normalizing CBF responses during adaptation to cortical hypoperfusion.

**Figure 8.**
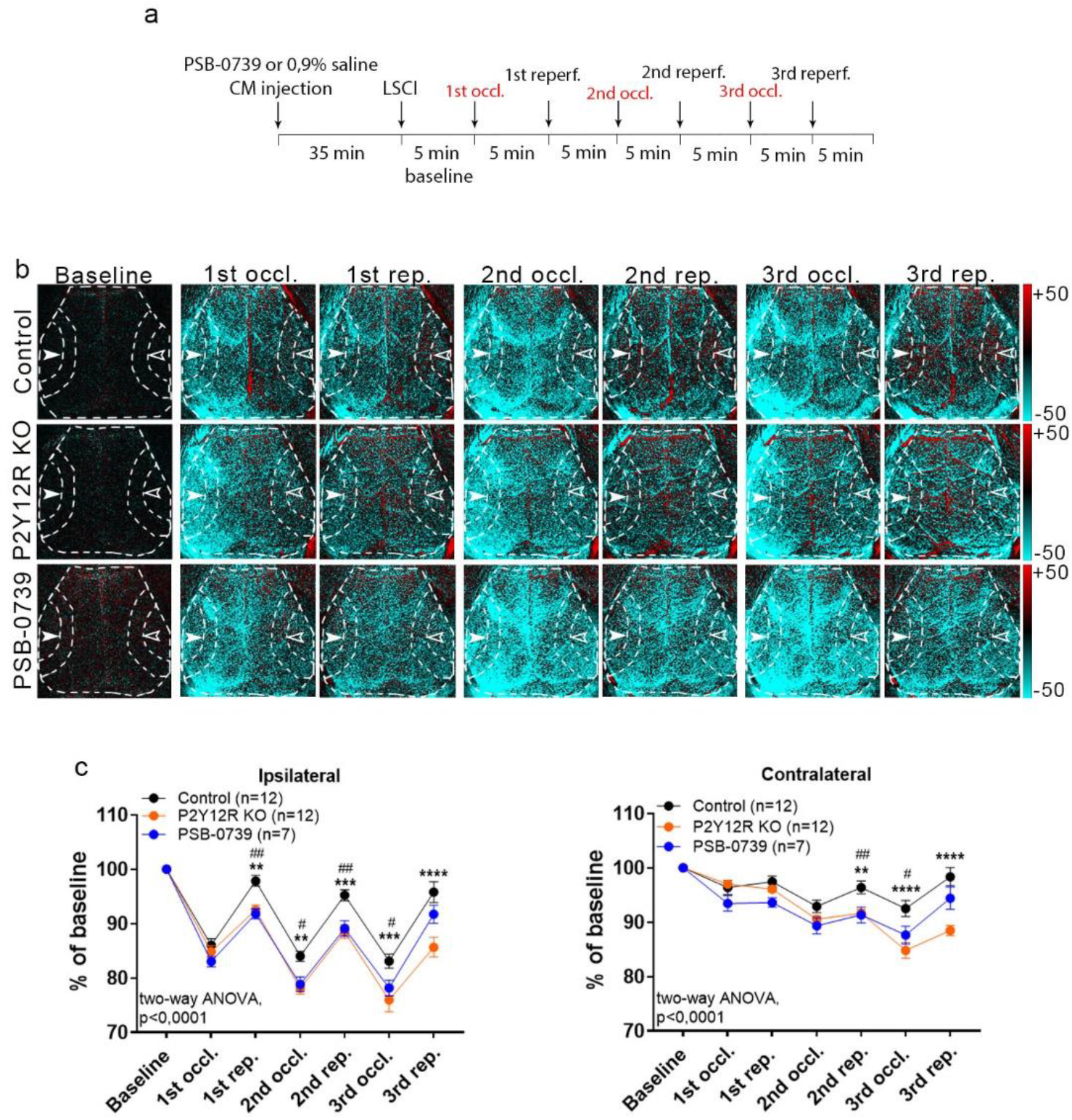
Microglial actions on cerebral blood flow require P2Y12R signaling. **a)** Outline of the experimental 3x CCAo protocol. CCAo was performed by using a silk thread. **b)** Representative difference images show altered perfusion in both hemispheres in response to 3x CCAo in P2Y12R KO and PSB-0739-injected mice compared to controls. Dashed lines show the MCA2 area both in the ipsilateral (white arrowheads)-, and in the contralateral hemisphere (unfilled arrowheads) corresponding to the quantitative analysis shown in panel c. **c)** A significant impairment in adaptation to hypoperfusion is seen both in the ipsilateral and contralateral hemispheres of P2Y12R KO mice and PSB-0739 injected mice compared to controls. n=12-12-7 mice. *p control vs P2Y12R KO mice, ^#^p control vs PSB-0739 injected mice, two-way ANOVA followed by Tukey’s multiple comp. test (c).

## Discussion

Here we identify microglia as a novel cell type regulating blood flow in the brain. Using three different experimental models, we show that the presence of functional microglia is essential to maintain optimal CBF responses to physiological neuronal activity, hypercapnia and during cerebrovascular adaptation to hypoperfusion. These actions are dependent on microglial P2Y12R signaling, clearly discriminating microglial responses from those mediated by PVMs or other brain macrophages ^52^.

While microglia produce several vasoactive or inflammatory mediators, including IL-1β, TNFα, NO, PGE2 or ROS ^4^ that may modulate cerebral perfusion ^11, 24^, the potential contribution of microglia to CBF has been largely neglected to date. Instead, research has focused on the role of microglia in BBB function, extravasation of leukocytes and angiogenesis from embryonic stages into adulthood ^10^. Because microglial cell bodies are located in the brain parenchyma, while the endothelial basal lamina is surrounded by a second, glial basement membrane ^53^, we first asked whether a direct contact between microglia and endothelial cells exists in the adult neocortex. We found microglia dynamically contacting different levels of the vascular tree *in vivo* and establish direct, purinergic contacts with endothelial cells, peri-arterial smooth muscle cells, pericytes and astrocytes in both the mouse and the human brain, which regulate blood flow. These observations suggested that purinergic mediators, such as ADP may be released from NVU cells to recruit P2Y12R-positive microglial processes during vascular adaptation responses or perfusion changes, even under physiological conditions.

Microglia depletion did not cause major anatomical changes in the NVU as also seen earlier ^21, 34^, confirmed by our [^99m^Tc]-HMPAO SPECT and [^18^F]-FDG PET measurements, which are widely used non-invasive methods to assess regional CBF and glucose metabolism changes, respectively ^32, 33^. To investigate whether microglia could influence CBF responses to physiological neuronal activity, we turned to the widely used whisker stimulation model. Neurovascular coupling is a dynamic functional change in CBF in response to local neuronal activity, which involves different cell types within the NVU, including astrocytes, vascular smooth muscle cells, pericytes and endothelial cells ^24, 25^. However, a role for microglia has not been previously established. During functional hyperemia, dilation of arterioles propagates at high speed in a retrograde direction to upstream arteries, including branches of pial arteries, with both arteriolar and capillary dilation playing a role in increased O_2_ delivery ^25^. Our LSCI studies revealed significantly smaller CBF response to whisker stimulation in the barrel cortex in the absence of microglia, or microglial P2Y12R, which was not explained by altered neuronal responses in the barrel cortex as assessed by *in vivo* electrophysiology or two-photon calcium imaging. While neuronal activity during hypercapnia was not different between control, P2Y12R KO and microglia-depleted mice either, it remains to be investigated whether microglia-dependent effects may also influence CBF through changing baseline activity of neurons that control blood flow in the brain ^25,23,54^. Importantly, fUS measurements confirmed microglia-dependent effects in CBF responses to somatosensory stimulation.

We next tested whether microglia-mediated mechanisms influence vascular responses to hypercapnia. Hypercapnia induces vasodilation via complex actions that involve NO release from the endothelium, activation of smooth muscle cells, pericytes, release of astrocytic prostaglandin E2 and other processes ^28, 39–42^. Importantly, while perivascular microglial processes rapidly responded to hypercapnia-induced vasodilation, the absence of microglia markedly inhibited increases of CBF (as demonstrated independently by both LSCI and laser Doppler flowmetry) and vasodilation (as shown by *in vivo* two-photon imaging). This was independent of arterial blood pH, pO_2_ and pCO_2_ levels, which were not different in microglia-depleted mice. Surprisingly, we found that absence of microglia reduced brain pH, while microglia rapidly produced adenosine in response to hypercapnia. These observations are also supported by recent findings showing that microglia represent a key source of adenosine in the brain to modulate neuronal responses at synapses ^54^. Hypercapnia drives vasodilation mainly via reduced extracellular pH, which is a major regulator of cerebrovascular reactivity and acts directly on cerebrovascular smooth muscle cells to cause relaxation, mediating the effects of increased CO_2_ levels ^41^. Microglial P2Y12 receptor-mediated Ca^2+^ signalling, migration and cytokine production are also pH-dependent ^55, 56^. Because adenosine is a potent vasodilator in the cerebral circulation ^57^, we suggest that lower brain pH in the absence of microglia may partially compensate for the loss of microglial vasoactive mediators with a net effect of reduced vasodilation during different vascular adaptation responses. It should be investigated in future studies whether microglial loss or dysfunction could induce compensatory actions in other NVU cells, such as promoting adenosine production by astrocytes^58^.

We then asked whether microglia sense and respond to cerebral hypoperfusion. Cerebral blood flow is controlled by feed-forward and feed-back mechanisms that maintain or re-establish optimal oxygen and nutrient supply of neurons in case disturbances of the cardiovascular system occur ^2^. Adaptation to hypoperfusion requires vasodilation^59^. Unilateral CCAo is an established model of cerebrovascular adaptation to hypoperfusion, which is mediated primarily by the activation of feed-back pathways through the collateral circulation^47^, while it does not induce neuronal death or BBB injury in rodents^47, 48^. Since the cell types in the NVU contacted by microglia regulate CBF ^24, 25, 29^, we argued that microglia primed by hypoperfusion during the first occlusion would interfere with subsequent vascular adaptation responses and hence elimination of microglia may alter CBF after repeated CCAo. As supported by previous results showing that microglial process responses around microvessels change proportionally to the level of CBF reduction ^22^, we found that microglial processes rapidly respond to CCAo. Importantly, absence of microglia and both genetic and pharmacological blockade of microglial P2Y12R resulted in impaired adaptation to hypoperfusion during repeated CCAo. We also found impaired CBF adaptation responses in CX3CR1^GFP/+^ mice. While CX3CL1 (fractalkine) is synthesized as a membrane protein by endothelial cells mediating leukocyte adhesion or chemotactic activity upon cleavage via acting on CX3CR1^60^, interactions between neuronal CX3CL1 and microglial CX3CR1 are widely recognized to control microglial activity ^50, 61^. These complex interactions may explain the controversial phenotype of CX3CR1^GFP/GFP^ mice on CBF changes during repeated CCAo, which strain is widely used to study microglia-mediated effects in the brain ^49^. Interestingly, V249 and T280 polymorphisms in the CX3CR1 gene show a remarkable association with an increased total arterial blood volume in the human brain ^62^, although this effect has not been functionally linked with microglial actions previously. Importantly, while a population of PVMs was also found to be CX3CR1-positive, both the effect of PVM depletion by clodronate on CBF in the repeated CCAo paradigm, and the reduced hypercapnia-induced vasodilation seen in P2Y12R KO × CX3CR1^GFP/+^ (double transgenic) mice confirmed the contribution of microglia to purinergic vascular adaptation responses and strengthens their different roles compared to P2Y12R-negative PVMs^23, 37^.

In most repeated CCAo experiments, the CCA was pulled away by a silk thread resulting in a reproducible, app. 20% reduction in cortical blood flow in control mice, allowing a gradual impairment in CBF recovery to be observed in response to microglia manipulation toward the third CCA occlusion and reperfusion. In studies investigating brain pH and DC potential, CCAo was performed by a mechanical occluder, which resulted in a larger (over 40%) and more abrupt CBF reduction, unexpectedly triggering SD, a large acidic pH shift and progressive oligaemia in the absence of microglia. Interestingly, we have previously reported altered extracellular potassium homeostasis after microglia depletion *in vivo*, with a lower initial SD threshold in the intact brain, but impaired SD induction after repeated SDs, or during hypoperfusion due to cerebral ischemia *^21, 63^*. In the presented study, microglia depleted mice immediately showed a robust, 80% drop in CBF during the first CCAo, followed by sustained impairment in CBF recovery, which was also seen in the milder CCAo model upon different microglia manipulations. Thus, these results collectively suggest that impaired CBF adaptation in the hypoperfused brain is a common phenomenon in the absence of functional microglia, which may be further augmented by SD-associated spreading oligaemia in case the CBF reduction is sufficient to trigger SD ^51^.

The importance of ATP signaling in the vasculature has been demonstrated under both homeostatic and pathological conditions ^27^. Microglial processes are recruited to sites of ATP release via P2Y12 receptors, which primarily sense ADP produced by ATP hydrolysis or cleavage by NTPDase1 expressed on the microglial membrane among other cells^23 7, 9^. Our electron tomography studies revealed an accumulation of P2Y12R on microglial processes contacting endothelial cells in the vicinity of endothelial mitochondria, where ATP release may recruit microglial processes to the vasculature in response to CBF changes ^27^. Similar interactions are seen at somatic purinergic junctions ^23^, through which microglia sense neuronal mitochondrial activity and modulate neuronal responses via purinergic signaling. ATP derived from astrocytes is also known to constrict vascular smooth muscle cells and regulate blood flow ^25^. Importantly, our HPLC studies demonstrated that endothelial cells and astrocytes release different purinergic metabolites in response to hypoxia and hypercapnia, both of which occur during hypoperfusion. While we found rapid alterations in microglia-endothelium and microglia-astrocyte interactions after CCAo and hypercapnia, hypoxia and hypercapnia also triggered different purinergic responses in microglia. Thus, cell- and stimulus-specific production of vasoactive metabolites may provide means for different vascular adaptation responses, during which microglia may alter CBF via actions on different cell types in the NVU or protect against mild hypoxia-induced vascular leakage ^64^. Functionally distinct responses in given microdomains may rely on the functional autonomy of calcium signaling in microglial processes ^65^.

### Clinical significance

We believe that the implication of these data is far-reaching. Microglial activity is altered in brain diseases, while microglial inflammatory mediators are important regulators of vascular permeability and CBF. Interestingly, common brain pathologies such as Alzheimer’s disease, Lewy body dementia, idiopathic Parkinson’s disease, chronic hypoperfusion, or amyloid angiopathy are associated with impaired CBF responses and/or neurovascular coupling often preceding the appearance of symptoms ^4, 24, 25, 29^. These changes are generally attributed to impaired neuronal, astrocyte, endothelial or pericyte responses. Based on our data, altered microglial activity or dysfunction of microglia could also contribute to disease pathophysiology by regulating cerebral perfusion directly via endothelial cells or through other cell types in the NVU. For example, in frontotemporal dementia, resting CBF is reduced in the frontal, parietal, and temporal cortex of presymptomatic patients, an effect independent of atrophy ^66^. Homozygous missense mutations of TREM2 (a microglial receptor) are linked with increased risk of dementia, while Trem2 p.T66M knock-in mice display an age-dependent reduction in microglial activity, cerebral blood flow and brain glucose metabolism ^67^. While the repeated CCAo model is useful to study how hypoperfusion-primed microglia modulate CBF, the present findings also have substantial pathophysiological relevance: in patients with risk factors for stroke, carotid stenosis, aneurysm, hypertension, chronic vascular inflammation or TIA, altered microglial activity due to perfusion changes or other factors may impact on clinical outcome after acute vascular events, or influence chronic neurodegeneration seen in AD or vascular dementia merely via adjusting cerebral perfusion or adaptation to hypoperfusion. As such, microglia could also contribute to ischemic preconditioning, vasospasm after subarachnoid haemorrhage or the „no reflow phenomenon” after cerebral ischemia ^68^, while microglial surveillance is likely to be disturbed during hypoxia or ischemia, as evidenced in the developing brain ^22, 69^. In conclusion, our data demonstrate that microglia should be considered as an important modulatory cell type involved in physiological and pathological alterations of cerebral blood flow and understanding their actions may facilitate the discovery of novel treatment opportunities in common neurological disorders.

## Materials and Methods

### Mice

Experiments were carried out on 11-17 weeks old C57BL/6J (RRID:IMSR_JAX:000664), P2Y12R^−/−^ (B6;129-P2ry12^tm1Dgen^/H P2Y12R KO), CX3CR1^GFP/+^ (RRID:IMSR_JAX:005582), CX3CR1^GFP/+^/P2Y12^−/−^, CX3CR1^GFP/GFP^, CX3CR1^tdTomato^, and Thy1-GCaMP6s (C57BL/6J-Tg(Thy1-GCaMP6s)GP4.12Dkim/J, RRID:IMSR_JAX:025776)^70^ mice (all on C57BL/6J background). Mice were kept in a 12h dark/light cycle environment, under controlled temperature and humidity with food and water ad libitum. All experimental procedures were in accordance with the guidelines set by the European Communities Council Directive (86/609 EEC) and the Hungarian Act of Animal Care and Experimentation (1998; XXVIII, Sect. 243/1998), approved by the Animal Care and Experimentation Committee of the Institute of Experimental Medicine and the Government Office of Pest County Department of Food Chain Safety, Veterinary Office, Plant Protection and Soil Conservation Budapest, Hungary under the numbers PE/EA/1021-7/2019, PE/EA/673-7/2019 and Department of Food Chain Safety and Animal Health Directorate of Csongrád County, Hungary. Experiments were performed according to the EU Directive 2010/63/EU on the protection of animals used for scientific purposes, and reported in compliance with the ARRIVE guidelines.

### Generation and characterization of CX3CR1^tdTomato^ microglia reporter mice

CX3CR1^tdTomato^ mice were generated by crossing tamoxifen-inducible CX3CR1^CreERT2^ mice (B6.129P2(C)-CX3CR1^tm2.1(cre/ERT2)Jung/J^, RRID:IMSR_JAX:020940 ^71^) with a cre-dependent tdTomato-expressing line (B6;129S6-Gt(ROSA)26Sortm9(CAG-tdTomato)Hze/J, RRID:IMSR_JAX:007905). To induce tdTomato expression in microglia, Cre recombinase activity was induced by two intraperitoneal injections of tamoxifen (TMX, 2mg/100μl, #T5648, Sigma, dissolved in corn oil), 48h apart in 3-4 weeks old male mice, shortly after weaning. Four weeks after TMX induction, virtually 100% of tdTomato expressing cells were identified as microglia, as assessed by P2Y12 and Iba1 immunostaining - tdTomato-positive cell bodies and processes were analyzed in the somatosensory cortex on high-resolution CLSM stacks (CFI Plan Apochromat VC 60XH oil immersion objective, 0.1μm/pixel, Z-step: 2 μm) with homogenous sampling (Supplementary Fig.1h).

### *In vivo* two-photon imaging

CX3CR1^GFP/+^, CX3CR1^tdTomato^, CX3CR1^GFP/+^ xP2Y12^−/−^ or Thy1-GCaMP6s mice were anaesthesized using 1,8% isoflurane or fentanyl (0.05mg/kg) and a 3 mm diameter cranial window was opened on the left hemisphere, above the primary somatosensory or the barrel cortex without damaging the dura mater. After removal of the skull bone, double circular glass coverslip was fixed with 3M™ Vetbond™. Above the dual coverslip, a custom-made metal headpiece (Femtonics Ltd., Budapest, Hungary) was fixed with dental cement. Three weeks after cranial window surgery, microglia-vascular interactions in response to 3x CCAo or hypercapnia (ketamine-medetomidine anesthesia, i.p. 30mg/kg-0,1mg/kg), and neuronal [Ca^2+^]_i_ in response to whisker stimulations (in ketamine-medetomidine anaesthesia) were imaged in body-temperature controlled animals. To image microglia vascular interaction, blood vessels were labeled either with Rhodamine B-Dextran (70.000 mol wt, neutral, #D1841, Molecular Probes) or with FITC-Dextran (70.000 mol wt, #53471, Sigma) injected into the retro-orbital sinus or into to the tail vein. Two-photon imaging was performed with a Femto2D-DualScanhead microscope (Femtonics Ltd., Budapest, Hungary) coupled with Chameleon Discovery laser (Coherent, Santa Clara, USA). The measurements were done either with the 920 nm tunable or with 1040 nm fixed wavelength laser excitation to simultaneously record GFP and Rhodamine B-Dextran signal as well as tdTomato and FITC-Dextran signal *in vivo* in real time. For microglia process motility measurements in response to 3x CCAo, the galvo-scanning light path with 16x water immersion objective (Nikon CFI75 LWD 16x W, NA 0.8) was used to acquire 4 images Z-stacks with 8.5 μm step size, 150-200 μm below the dura, at 500 × 500 pixel resolution. For measuring vascular responses to hypercapnia, a mild ketamine-medetomidine sedation (i.p. 30 mg/kg-0,1 mg/kg) was used. After obtaining 1 min baseline, a 2 min hypercapnic episode (inhaling a 10 % CO_2_-containing air mixture) and 1 min post-hypercapnic period were imaged with the resonant light path at 32.7521 Hz. For *in vivo* [Ca^2+^]_i_ imaging in control and microglia depleted Thy1-GCaMP6s mice, three weeks after cranial window surgery the right whiskers were stimulated with a bending actuator (#PL112-PL140 PICMA Bender) connected to a piezo amplifier (#E-650 Amplifier, Physik Instrumente (PI) GmbH, Karlsruhe, Germany), and neuronal [Ca^2+^]_i_ transients were imaged in the left barrel cortex, at 180-250 μm depth below the dura, under ketamine-medetomidine sedation. The stimulation protocol consisted of 5Hz square pulses for 15 seconds that was repeated 6 times with 40 seconds intervals. Measurements to detect the GCaMP6s signal were performed at 920 nm wavelength, using the resonant scanner at 32.48 Hz and a Nikon 16x water immersion objective. Data acquisition was performed by MESc software (v.3.5.6.9395SLE, Femtonics Ltd.) and data analysed by MES software (v.5.3560, Femtonics Ltd.).

### Tissue processing and immunostaining

Under terminal (ketamine-xylazine) anaesthesia mice were transcardially perfused with 4% paraformaldehyde (PFA) and brains were dissected. Brain samples were post-fixed, cryoprotected for 24h and 25 μm thick coronal sections were cut, using a sledge microtome (Leica, Germany). Immunostaining was performed on free-floating brain sections, blocked with 5% normal donkey serum (Jackson ImmunoResearch Europe Ltd, Ely, Cambridgeshire, UK). The following primary antibodies were used: rabbit anti-P2Y12R (1:500, #55043AS AnaSpec), chicken anti-GFP-tag (1:500, #A10262 Invitrogen), rat anti-CD206 (1:200, #MCA2235, AbD Serotec) and biotinylated tomato lectin (1:100, #B-1175, Vectorlabs). After washing, sections were incubated with the corresponding secondary antibodies: donkey-anti chicken A488 (1:500, #703-546-155, Jackson Immunoresearch), streptavidin DyL405 (1:500, #016-470-084, Jackson ImmunoResearch), donkey anti-rabbit A647 (1:500, #711-605-152, Jackson ImmunoResearch) and donkey anti-rat A594 (1:500, #A21209, Invitrogen). Slices were mounted with Fluoromount-G (SouthernBiotech) or Aqua-Poly/Mount (Polysciences). For high resolution confocal laser scanning microscopy (CLSM) and electron microscopic assessments, 50 μm thick vibratome sections were washed in PB and TBS, followed by blocking with 1% human serum albumin (HSA). Sections were then incubated in different mixtures of primary antibodies: rabbit anti-P2Y12R (1:500, #55043AS AnaSpec), chicken anti-GFAP (1:500, #173 006 Synaptic Systems), goat anti-PDGFR-β (1:500, #AF1042 R&D Systems), rat anti-CD206 (1:200, #MCA2235, AbD Serotec), rat anti-PECAM-1 (1:500, #102 501, BioLegend), mouse anti-αSMA (1:250, #ab7817, Abcam), guinea pig anti-Aquaporin-4 (1:500, #429 004, Synaptic Systems), mouse anti-TOM20 (1:500, #H00009804-M01, Abnova), guinea pig anti-Iba1 (1:500, #234 004, Synaptic Systems), mouse anti-Kv2.1 (1:500, #75-014, NeuroMab) and biotinylated tomato lectin (1:100, #B-1175, Vectorlabs). After washing in TBS, sections were incubated in the corresponding mixtures of secondary antibodies: donkey anti-chicken DyLight405 (1:500, #703-474-155, Jackson ImmunoResearch), donkey-anti chicken A488 (1:500, #703-546-155, Jackson ImmunoResearch), donkey anti-chicken A647 (1:500, #703-606-155, Jackson ImmunoResearch), donkey anti-rabbit A647 (1:500, #711-605-152, Jackson ImmunoResearch), donkey anti-rabbit A488 (1:500, #A21206, Invitrogen), donkey anti-rat A594 (1:500, #A21209, Invitrogen), donkey anti-rat A647 (1:500, #712-606-153, Jackson ImmunoResearch), donkey anti-mouse A594 (1:500, #A21203, Invitrogen), donkey anti-mouse A647 (1:500, #715-605-150, Jackson ImmunoResearch), donkey anti-guinea pig DyLight405 (1:500, #706-476-148, Jackson ImmunoResearch), donkey anti-guinea pig A594 (1:500, #706-586-148, Jackson ImmunoResearch), donkey anti-guinea pig A647 (1:500, #706-606-148, Jackson ImmunoResearch), streptavidin DyL405 (1:500, #016-470-084, Jackson ImmunoResearch), streptavidin A594 (1:500, #S11227, Invitrogen). Incubation was followed by washing in TBS and PB, then sections were mounted on glass slides with Aqua-Poly/Mount (Polysciences). Immunofluorescence was analyzed using a Nikon Eclipse Ti-E inverted microscope (Nikon Instruments Europe B.V., Amsterdam, The Netherlands), with a CFI Plan Apochromat VC 60X oil immersion objective (NA 1.4) or a Plan Apochromat VC 20X objective (NA 0.75) and an A1R laser confocal system. The following lasers were used: 405, 488, 561 and 647 nm (CVI Melles Griot). Scanning was done in line serial mode. Image stacks were obtained with NIS-Elements AR 5.00.00 software.

### Pre-embedding immunoelectron microscopy

After extensive washes in PB and TBS (pH 7.4), vibratome sections were blocked in 1 % HSA. Then, they were incubated with rabbit anti-P2Y12R (1:500, #55043AS AnaSpec) alone or mixed with mouse anti-GFAP (1:1000, #G3893 Sigma) in TBS for 2-3 days. After several washes, sections were incubated in blocking solution (Gel-BS) containing 0.2% cold water fish skin gelatine and 0.5 % HSA for 1 h. Next, sections were incubated with 1.4 nm gold-conjugated goat anti-rabbit Fab-fragment (1:200, #2004 Nanoprobes) alone or mixed with biotinylated donkey anti-mouse (1:500, #715-065-150 Jackson Immunoresearch) antibodies diluted in Gel-BS overnight. After extensive washes, sections were treated with 2 % glutaraldehyde for 15 min to fix the gold particles into the tissue. For the combined immunogold-immunoperoxidease reactions, this was followed by an incubation in avidin–biotinylated horseradish peroxidase complex (Vectastain Elite ABC kit; 1:300; Vector Laboratories) for 3 h at room temperature (RT) or overnight at 4°C. The immunoperoxidase reaction was developed using 3,3-diaminobenzidine (DAB; Sigma-Aldrich) as chromogen. To develop the immunogold reaction, sections were incubated in silver enhancement solution (SE-EM; Aurion) for 40-60 min at RT. The sections were then treated with 0.5 % OsO_4_ in PB, at RT, dehydrated in ascending alcohol series and in acetonitrile, and were embedded in Durcupan (ACM; Fluka). During dehydration, sections were treated with 1 % uranyl acetate in 70 % ethanol for 20 min. For electron microscopic analysis, tissue samples from the somatosensory cortex (S1) were glued onto Durcupan blocks. Consecutive 70 nm-thick (for conventional electron microscopic analysis) or 150 nm-thick (for electron tomography) sections were cut using an ultramicrotome (Leica EM UC6) and picked up on Formvar-coated single-slot grids. Ultrathin sections for conventional electron microscopic analysis were examined in a Hitachi H-7100 electron microscope equipped with a Veleta CCD camera (Olympus Soft Imaging Solutions, Germany). 150 nm-thick electron tomography sections were examined in FEI Tecnai Spirit G2 BioTwin TEM equipped with an Eagle 4k camera.

### Electron tomography, analysis

Before electron tomography, serial sections on single-slot copper grids were photographed with a Hitachi H-7100 electron microscope and a Veleta CCD camera. Serial sections were examined at lower magnification, and P2Y12R-positive microglial processes contacting the vasculature were selected. After this, grids were put on drops of 10 % HSA in TBS for 10 minutes, dipped in distilled water (DW), put on drops of 10 nm gold conjugated Protein-A (Cytodiagnostics #AC-10-05) in DW (1:3), and washed in DW. Electron tomography was performed using a Tecnai T12 BioTwin electron microscope equipped with a computer-controlled precision stage (CompuStage, FEI). Acquisition was controlled via the Xplore3D software (FEI). Regions of interest were pre-illuminated for 4-6 minutes to prevent further shrinkage. Dual-axis tilt series were collected at 2 degree increment steps between −65 and +65 degrees at 120 kV acceleration voltage and 23000x magnification with −1.6 – −2 μm objective lens defocus. Reconstruction was performed using the IMOD software package ^72^. Isotropic voxel size was 0.49 nm in the reconstructed volumes. After combining the reconstructed tomograms from the two axes, the nonlinear anisotropic diffusion (NAD) filtering algorithm was applied to the volumes. Segmentation of different profiles has been performed on the virtual sections using the 3Dmod software.

### *Post mortem* human brain samples

*Post mortem* human brain tissue was obtained from one 60-years-old female, one 73-years-old male and one 27-years-old male patient without any known neurological disease as also confirmed by neuropathological examination (ETT TUKEB 31443/2011/EKU [518/PI/11]). Informed consent was obtained for the use of brain tissue and for access to medical records for research purposes. Tissue was obtained and used in a manner compliant with the Declaration of Helsinki. Brains of patients who died in non-neurological diseases were removed 4-5 h after death (Supplementary Table 1). The internal carotid and the vertebral arteries were cannulated, and the brain was perfused first with heparin containing physiological saline, followed by a fixative solution containing 4% PFA, 0.05% glutaraldehyde and 0.2% picric acid (vol/vol) in PB. The hippocampus was removed from the brain after perfusion, and was postfixed overnight in the same fixative solution, except for glutaraldehyde, which was excluded. Blocks were dissected, and 50 μm thick sections were prepared on a vibratome (VT1200S, Leica, Germany).

### Laser Speckle Contrast Imaging (LSCI)

CBF was measured by a PeriCam PSI High Resolution LSCI system (Perimed AB, Järfälla-Stockholm, Sweden). Before CBF measurements, the head of the mouse was secured in a stereotaxic head holder, and after a midline incision, the skull was exposed by retracting the scalp. Imaging was performed through the intact skull bone. The cerebrocortical microcirculation was imaged at 21 frames/sec frequency in a 10×10 mm field of view. Perfusion responses were expressed as a percentage of baseline CBF. Uniformly, a 1 min long baseline was set in all experiments, registered at the beginning of the measurements. Three adjacent ROIs were placed (denoted as MCA1-3) over the middle cerebral artery (MCA) territory both to the ipsilateral and to the contralateral hemispheres to assess microglia-mediated effects on gradual perfusion changes ranging from the MCA core region to the midline. CCA occlusion experiments were performed under ketamine-xylazine (i.p. 100mg/kg - 10mg/kg) anaesthesia. The whisker stimulation protocol was performed under mild ketamine-medetomidine (i.p. 30mg/kg – 0.1mg/kg) sedation ^73^ with ROIs placed over the contralateral barrel cortex. The hypercapnic challenge was done under mild ketamine-medetomidine (i.p. 30mg/kg – 0.1mg/kg) sedation and ROIs were placed over the left and right hemispheres excluding venous sinuses.

### Cisterna magna injection for drug delivery into the brain

To block P2Y12 receptor (P2Y12R)-mediated microglial actions, a P2Y12R antagonist, PSB-0739 (dissolved in 0.9% saline, 40 mg/kg in 5 μl volume, #3983, Bio-Techne Corp., Minneapolis, USA) was injected into the cisterna magna 35 min prior imaging, while vehicle (0.9% saline) injection was used as control. Cisterna magna injections were done under 1-1.5% isoflurane anaesthesia.

### SPECT and PET imaging

Single photon emission computed tomography (SPECT) and positron emission tomography (PET) studies were carried out on mice anaesthetized with 2% isoflurane ^32, 33^. SPECT measurements were performed using the [^99m^Tc]-HMPAO ligand (Hexamethylpropylene amine oxime, Medi-Radiopharma Co Ltd., Hungary). The acquisition started 3 minutes after the i.v. injection of the radiotracer via the tail vein (injected activity: 99.22 ± 9.33 MBq). The measurements were performed on a NanoSPECT/CT PLUS device (Mediso Ltd, Hungary) equipped with multi-pinhole mouse collimators. Measurements were reconstructed with 0.25 mm isovoxels and the results were quantified as units of radioactivity (MBq/ml). After SPECT acquisition, [^18^F]-FDG (2-deoxy-2-(18F)fluoro-D-glucose) PET measurements were performed. PET acquisition started 20 minutes after i.v. [^18^F]-FDG injection (Pozitron-Diagnosztika Ltd, Hungary; injected activity: 12.05 ± 1.93 MBq) with an acquisition time of 10 minutes using a microPET P4 (Concorde Microsystems Inc, USA). A maximum a posteriori algorithm was used to reconstruct the data with 0.3 mm isovoxels. After reconstruction, manual coregistration and atlas-based region of interest (ROI) measurements were done using VivoQuant software (InviCRO, USA) in the cerebellum, cerebral cortex and the whole brain. For microglia depleted and control groups, mean [18F]-FDG and [99mTc]-HMPAO standardized uptake values (SUV) were analyzed by using two-way ANOVA followed by Sidak’s post-hoc test (GraphPad Prism 7.0) and a permutation t-test in R 3.5.1 (R Foundation for statistical computing, Austria).

### Whisker stimulation protocol

Whisker stimulation was performed manually and electromechanically (with a bending actuator #PL112-PL140, PICMA, Bender connected to a piezo amplifier #E-650 Amplifier, Physik Instrumente (PI) GmbH, Karlsruhe, Germany). For manual stimulation an earpick was used (at 4-5 Hz frequency) according to the following protocol: left whiskers were stimulated for 30 sec, repeated 6 times, 60 sec apart. During electromechanically controlled stimulation (5 Hz) whiskers were stimulated for 15 sec, repeated 10 times with 40 sec intervals. Stimulation-evoked CBF responses in the contralateral barrel cortex were recorded. CBF measurements were carried out under ketamine-medetomidine sedation (30 mg/kg - 0.1 mg/kg dissolved in 0.9% saline, i.p.). All coupling experiments performed were time-matched from the time of anesthetic injection to ensure comparable results across different experiments.

### Functional ultrasound (fUS)

The acquisition was done with a 15 MHz probe of 128 elements (Vermon SA, France) connected to a prototype ultrafast research ultrasound scanner (hardware and software functionally analogous to the Iconeus One system, Iconeus, Paris, France). Recordings were performed through the skull while the animal was anaesthetised with ketamine-medetomidine (i.p. 30mg/kg - 0.1mg/kg). The head of the animal was shaven and fixed into a stereotactic frame. The probe was positioned using a built-in software based registration to the 3D Allen Brain Atlas (2015 Allen Institute for Brain Science, Allen Brain Atlas API, available from: brain-map.org/api/index.html). The Doppler Images were obtained as described earlier^74^. 11 tilted planes were insonificating the medium at 5500 Hz pulse repetition frequency to compute one compounded image every 2 ms. Out of a block of 200 images a Power Doppler image was obtained by removing the 60 first modes of SVD decomposition to extract the blood signal^75^ from tissue clutter at a 2.5 Hz sampling rate. Acquisition started and ended with a 5 min baseline followed by 10 phases of 30 s manual stimulation of the whiskers^76^ with 1 min of resting in between. A fourth order polynomial detrending of the data was applied to remove drifts of baseline^77^.

### *In vivo* electrophysiology

Surgical procedures, microdrive construction and implantation have been described previously^78^. Briefly, custom-built microdrives with eight nichrome tetrodes (diameter, 12.7 μm, Sandvik, Sandviken, Sweden) and a 50-μm core optic fiber (outer diameter, 65± 2 μm, Laser Components GmbH, Olching, Germany) were implanted into the right barrel cortex AP: −1.4; ML: 3.0 DV 0.75−2.0 mm. Although photostimulation was not applied here, the optic fiber is part of our typical drive design as it also provides mechanical support for the tetrodes. The microdrive contained a moveable shuttle allowing more precise targeting. The custom-built microdrives were implanted under deep anaesthesia using an intraperitoneal injection of ketamine - xylazine (125 mg/kg - 25 mg/kg in 0.9% NaCl). Lidocain spray was used on the skin of the scalp to achieve local analgesia. The skin was incised, the skull was cleaned and leveled, and a cranial window was opened above the target area. Fluorescent dye (DiI, #LSV22885, Invitrogen) was applied on the tip of the tetrodes for later histological localization. Implants were secured by dentil adhesive (C&B Metabond, Parkell, Edgewood, NY, USA) and acrylic resin (Jet Denture, Lang Dental, Wheeling, USA). Buprenorphine was used for post-operative analgesia (Bupaq, 0.3 mg/ml, Richter Pharma AG, Wels, Austria). The stereotaxical surgery was followed by a 3 day-long resting period. During electrophysiological recordings animals were anesthetized using an intraperitoneal injection of a ketamine – medetomidine (3mg/kg – 0.1 mg/kg). The experiment was repeated twice or three times, with a two days gap between sessions. Tetrodes were lowered (40-120 μm based on the estimated electrode positions and the presence of single units) between recording sessions to collect neuronal activity from different dorsoventral position. Every session started with a 5min recording without stimulation, defined as basal activity. Automated whisker stimulation epochs lasted for 15 seconds with 5 Hz frequency, followed by a 40-second-long interstimulus period. During the stimulation, 5Hz frequency was generated with a TTL pulse generator, where the pulse duration and the interpulse interval was 0.1-0.1 second. In every 0.2 seconds the whisker stimulator moved forward, and after 0.1 second it passively moved backward to the starting position. This pattern resulted a bidirectional 10Hz stimulation. Stimulation was repeated 10 times. The entire protocol was repeated with the stimulator positioned close to the whiskers without touching them, to provide a sham stimulation condition that allowed us to exclude possible contaminations from electric noise from the stimulator circuit in our recordings. Next, manual whisker stimulation was applied (15 seconds stimulation with 40 seconds interstimulus period, repeated two times). Finally, changes in neuronal firing were measured during a 2 minutes long hypercapnic challenge, by inhalation of a 10% CO_2_ containing air mixture (21.1% O_2_ and N_2_ ad 100%) under normoxic conditions. After the last experiment, animals were terminally anesthetized and were transcardially perfused with 0.1 M PBS for 1 min, then with 4% PFA in PBS. After perfusion, brains were post-fixed and sections (50 μm coronal sections, Vibratome VT1200S, Leica) were imaged by fluorescence microscope (Nikon Eclipse Ni microscope, Nikon Instruments). Images were aligned to coronal sections of the Paxinos and Franklin atlas to accurately reconstruct the recording locations^79^. Data acquisition was conducted with an Open Ephys (open source data acquisition system, hiv4) board, synchronized with the electromechanical whisker stimulator through a pulse generator (PulsePal 1102, Sanworks)^80^. Data analysis was performed in Matlab R2016a (Mathwoks, Natick, US). Spike sorting was carried out using MClust 3.5 (A.D. Redish). Only neurons with isolation distance > 20 and L-ratio < 0.15 (a cluster quality measure based on Mahalanobis-distance^81^) were included.

### Induction of hypercapnia *in vivo*

Under mild ketamine-medetomidine (i.p. 30 mg/kg - 0.1 mg/kg) sedation, baseline cortical blood flow was recorded with LSCI for 5 minutes, then hypercapnia was induced by inhalation of a 10% CO_2_-containing air mixture (21.1% O_2_ and N_2_ ad 100%) for 2 minutes under normoxic conditions, followed by a 2 minutes long post-hypercapnic imaging period. In a group of control and microglia-depleted mice, before the hypercapnic challenge, 0.01 μg/g atipamezole (CP-Pharma, Revertor, 5mg/ml) was administered i.p. to withdraw α-2-agonistic effects of medetomidine. Three to five minutes were allowed to get the effect of atipamezole established, before recording baseline CBF and induction of hypercapnia.

### Blood gas analysis

Under ketamine-medetomidine anaesthesia (30mg/kg – 0.1mg/kg ± 0.1μg/g atipamezole), the left femoral artery was exposed and cannulated for arterial puncture. Arterial blood (45-75μl) was sampled to glass capillaries before and after 2min hypercapnic challenge (induced by inhalation of a 10% CO_2_ containing air mixture [21.1% O_2_ and N_2_ ad 100%] under normoxic conditions), and samples were measured with a blood gas analyzer (ABL90 FLEX PLUS, Radiometer Medical, Brønshøj, Denmark) to determine arterial blood gas tensions (pO_2_, pCO_2_) and pH.

### Repeated, transient CCA occlusion

Transient, repeated unilateral common carotid artery (CCA) occlusion was performed to induce hypoperfusion without causing ischemia or cellular injury to the brain. The CCA was temporarily pulled away with a silk suture for 5 min (except for the experiment presented in Fig.7c-f, as described under CBF, DC potential and brain pH measurements), followed by a 5 min-long reperfusion period. The protocol consisted of repeating these steps three times (3x CCAo) on anesthetised (ketamine-xylazine, i.p. 100mg/kg - 10mg/kg dissolved in 0.9% saline) mice. During CBF measurements, the core temperature of mice was maintained at 37 ± 0.5°C using a homeothermic blanket.

### Elimination of microglia or perivascular macrophages

C57BL/6J mice were fed a chow diet containing the CSF1R inhibitor, PLX5622 (Plexxikon Inc., 1200 mg PLX5622 in 1 kg chow) to eliminate microglia from the brain ^21^, or with control diet for 3 weeks. Perivascular macrophages (PVMs) were depleted by a single dose of clodronate-containing liposomes (70μg/mouse in 10μl volume, #F70101C-N-2, FormuMax Scientific, Inc.) injected into the left ventricle (ICV) as described earlier ^19^. Three days later, at maximal efficacy of depletion, laser speckle contrast imaging (LSCI) was carried out.

### Simultaneous measurement of CBF, DC potential and brain pH during repeated CCAo and hypercapnia

Mice were anesthetized with 1.5–2% isoflurane in N_2_O:O_2_ (3:2) and were allowed to breathe spontaneously through a head mask. Body temperature was maintained at 37°C with a servo-controlled heating pad (#507222 F, Harvard Apparatus, USA). Surgical preparations for CCAo were performed as described above. Subsequently, the animal was fixed in a stereotaxic frame, and a craniotomy (3-4 mm lateral from sagittal suture, −1 mm caudal from bregma) was created with a dental drill (ProLab Basic, Bien Air, Switzerland) over the somatosensory cortex on the right parietal bone. The dura mater in each craniotomy was left intact. Electrophysiological variables (DC potential, brain pH) and local CBF (by lased Doppler) were continuously monitored in the craniotomy. To this end, ion-sensitive microelectrodes were prepared according to Voipio and Kaila (1993) ^82^. In each experiment, a pH-sensitive microelectrode was lowered into the cortex with a micromanipulator, together with another glass capillary microelectrode (tip diameter = 20 μm) filled with saline to serve as reference. The tips of the two electrodes were positioned as near as possible. The reference electrode acquired slow cortical or DC potential. An Ag/AgCl electrode was implanted under the skin of the animal’s neck to be used as common ground. Microelectrodes were connected to a custom-made dual-channel high input impedance electrometer (including AD549LH, Analog Devices, Norwood, MA, USA) via Ag/AgCl leads. The voltage signal recorded by the reference electrode was subtracted from that of the pH-sensitive microelectrode by dedicated differential amplifiers and associated filter modules (NL106 and NL125, NeuroLog System, Digitimer Ltd., United Kingdom), which yielded potential variations related to changes in H^+^ ion concentration. The recorded signals were then forwarded to an analogue-to-digital converter (MP 150, Biopac Systems, Inc). Electric signals were continuously acquired at a sampling frequency of 1 kHz, displayed live, and stored using a personal computer equipped with the software AcqKnowledge 4.2.0 (Biopac Systems Inc., USA). Extracellular pH changes were expressed in mV to be translated into pH units offline, using least squares linear regression. In order to assess changes in local CBF, a laser-Doppler needle probe (Probe 403 connected to PeriFlux 5000; Perimed AB, Sweden) was positioned adjacent to the intra-cortical microelectrode at an angle (n = 14). The laser-Doppler flow (LDF) signal was digitized and displayed together with the DC potential and pH signals as described above (MP 150 and AcqKnowledge 4.2.0, Biopac Systems, Inc. USA). The completed preparations were enclosed in a Faraday cage. After the completion of the preparation, isoflurane was lowered to 0.5% and medetomidine was administered for the period of actual data acquisition (initiation: i.p. 0.5 mg/kg, repeated 5 min later for maintenance). After 15 min baseline acquisition, 2min hypercapnia was imposed by CO_2_-enriched gas inhalation (9.7 % CO_2_, 21 % O_2_ in N_2_, Messer, Hungary) at spontaneous respiration, which was repeated after a 5 min resting period. 10 min after the second hypercapnic challenge, transient repeated CCAo was performed as described above.

### Primary endothelial cells

Primary endothelial cultures were prepared from 6-8 weeks old C57BL/6J mouse brains as described originally by Deli et al.^83^, now performed with modifications according to Lenart et al.^84^. In brief, mouse forebrains were collected to PBS and the meninges were removed using sterile chromatography paper. The tissue was cut into small pieces by scalpels and was enzymatically digested in a mixture of Collagenase II (CLS2, 1mg/ml, #C6885, Sigma) and DNase I (0.025mg/ml, ~50U, #D4513, Sigma) in DMEM-F12 (#10-103-CV, Corning) for 55 min at 37C°. Using a 20% BSA (#A7906, Sigma, in DMEM-F12) gradient (1000x g, 20 min, three times), microvessels were separated from the myelin. The collected microvessels were further digested using a mixture of Collagenase/Dispase (1mg/ml, #11097113001, Sigma) and DNase I (0.038mg/ml, 75U, #D4513, Sigma) for 35 min at 37C°. Digested cerebral microvessels were washed three times with DMEM-F12, then seeded to Collagen type I (#354236, Corning) coated plates. During the first four days, puromycin ^85, 86^ (4μg/ml, #P7255, Sigma) selection was applied in the primary medium (15% PDS [#60-00-850, First Link] for seeding, 10% for cultivation, 1ng/ml bFGF [#F0291, Sigma], 100μg/ml heparin [#H3149, Sigma], 100x ITS [#41400045, Gibco], 4μg/ml puromycin in DMEM-F12) to selectively eliminate P-gp non-expressing cells. After reaching confluency in 5-6 days, the cells were passaged to Collagen type IV (100μg/ml, #C5533, Sigma) and fibronectin (25μg/ml, #F1141, Sigma) coated 48-well plates at a cell density of 15.000 cells/well, and used for *in vitro* hypoxia or hypercapnia experiments in P1.

### Primary astroglia and microglia cells

Primary cultures of astroglial cells were prepared from neonatal mouse brains, as described earlier^87^. In brief, meninges were removed from P0-P2 whole brains and tissues were chopped. The tissue pieces were digested with 0.05% w/v trypsin and 0.5mg/ml DNAse I (both from Sigma, #T4549, #DN25) in phosphate-buffered saline for 10min, at RT. Cells were then plated onto poly-L-lysine (#P1524, Sigma) coated plastic surfaces at a cell density of 3–4 × 10^5^ cell/cm^2^. The cultures were grown in Minimal Essential Medium (#21090022, ThermoFisher) supplemented with 10% fetal bovine serum (#FB-1090, BioSera), 4 mM glutamine (#G3126, Sigma), and 40 μg/ml gentamycin (Sandoz). The culture medium was changed twice a week. For the hypoxia/hypercapnia experiments, the primary cultures were passaged and plated in 1,5 × 10^5^ cell/cm^2^ density into poly-L-lysine coated 48 well plates and used within 96hrs. Astrocytes no older than 6-8 days in vitro were used. Primary microglia cells were isolated from astroglia/microglia mixed cultures derived from the whole brains of C57BL/6J newborn mouse pups. Microglia isolation was performed between days 21 and 28 of culture maintenance, by mild trypsinization^88^. For the *in vitro* hypoxia and hypercapnia experiments the isolated cells were seeded in a 1,5 × 10^5^ cell/cm^2^ density into poly-L-lysine coated 48 well plates and used within 48hrs.

### *In vitro* hypoxia and hypercapnia

Cytation 5 Cell Imaging Multi-Mode Reader (BioTek) equipped with O2/CO_2_ gas controllers was used to maintain 1% O2/5% CO2/94% N2 (Hypoxia) or 15% CO2/85% air (Hypercapnia) levels at 37°C. Endothelial or astroglial cells grown in 48 well plates to confluency, or microglia were placed into the reading chamber of the instrument for 5 (Hypercapnia) or 10 minutes (Hypoxia), after taking the lids off. In order to avoid medium-change induced release events, cell culture medium was replaced with 400ul complete fresh medium 16hrs before the onset of the experiments. To follow the build-up of hypoxia at the cellular level some cultures were loaded with 5uM Image-iT™ Green Hypoxia Reagent (#I14834, Invitrogen) for 30min at 37°C. The Hypoxia Reagent begins to fluoresce when atmospheric oxygen levels drop below 5%. Fluorescent images taken with Cytation5 (10x magnification) at 0/10minutes were analyzed with FIJI software (v1.53, NIH), measuring mean gray values in 10×10 pixel ROIs of n=50 individual cells from 3 independent experiments. Changes in medium pH during hypercapnia were measured by Phenol Red absorbance at 415 and 560 nm using the Cytation 5 Multi-Mode Reader (BioTek)^89^. Measurements were taken from 400 μL complete cell culture media in 48-well plates at 37 °C (n=10). The ratios of the 415 nm and 560 nm peaks were analyzed against a calibration curve obtained from 10mg/L phenol red and 10% FBS containing phosphate buffer solutions at different pH, in the range of pH=5,5-8. Changes in the intracellular pH during hypercapnia were determined by fluorescence intensity readings of pHrodo Green AM (#P35373, Invitrogen) labeled glial cells with Cytation 5 Multi-Mode Reader (n=4). Intracellular pH calibration was performed by incubating the pHrodo Green AM labeled cells in ACSF set to different pH values between pH=5,5-7,5 and supplemented with 10uM Nigericin and 10uM Valinomycin (#431, Nigericin; #3373 Valinomycin, Bio-Techne Corp., Minneapolis, USA), for 5 minutes.

### Quantification of nucleotides and nucleoside

Released concentrations of adenine nucleotides (ATP, ADP, AMP) and adenosine (Ado) from culture media were determined using HPLC method. The medium (400 μl) was transferred into a cold Eppendorf tube which contained (50 μl) of 0.1 M perchloric acid with 10 μM theophylline (as an internal standard) solution. The medium sample was centrifuged (at 3510 g for 10 min at 0-4°C) and the supernatant was kept at −20°C until analysis. Online Solid Phase Extraction coupled to the column-switching technique was applied to quantification of the nucleotide content of samples. HPLC separation was performed by Shimadzu LC-20 AD Analytical System using UV (Agilent 1100 VW set at 253 nm) detection. The phenyl-hexyl packed (7.5 × 2.1 mm) column was used for online sample enrichment and the separation was completed by coupling the analytical C-18 (150 × 2.1 mm) column. The flow rate of the mobile phases [“A” 10 mM potassium phosphate buffer with 0.25 mM EDTA and phase “B” contained additional components such as 0.45 mM octane sulphonyl acid sodium salt, 8% acetonitrile (v/v), 2% methanol (v/v), and the pH 5.55] was 350 or 450 μl/min, respectively in a step gradient application^90^. The sample enrichment flow rate of buffer “A” was 300 μl/min during 4 min and the total runtime was 55 min. Concentrations were calculated by a two-point calibration curve using internal standard method. The data (n=4 or 5 in each group) are expressed as nmol per mL. The Shapiro-Wilk normality test of the GraphPad Prism 8.4.3 statistical program confirmed the normal distribution of the data. Multiple t-test, corrected for multiple comparisons with the Holm-Sidak method was used for evaluation. The threshold for statistical significance was p <0.05.

### Quantitative analysis

All quantitative analyses were done in a blinded manner. For the measurements of microglial process coverage of endothelial surface or pericytes, microglial process coverage was measured on confocal Z-stacks acquired with a step size of 300 nm. On single-channel images, lectin-positive vessels were selected randomly. The surface of these vessels was calculated by measuring their circumference on every section multiplied by section thickness. The length of microglial process contacts was measured likewise. Continuous capillary segments (shorter than 6 μm) were also randomly chosen, and the presence of microglial process contacts was examined. All labeled and identified pericytes (PDGFRβ-positive) were counted when the whole cell body was located within the Z-stack. 3-dimensional reconstruction of CLSM and 2-photon imaging stacks was performed using the IMOD software package ^72^. TOM20 fluorescent intensity profiles were analyzed using a semi-automatic method (Supplementary Figure 1 d). Confocal stacks with triple immunofluorescent labeling (P2Y12R, TOM20 and Lectin) were collected. The image planes containing the largest diameter of longitudinal or cross-cut vessels were used to trace the outer membrane of endothelial cells based on the Lectin-labeling. This contour was then expanded and narrowed by 0.5 μm to get an extracellular and an intracellular line, respectively. The intensity of fluorescent labeling was analyzed along these lines (TOM20 intensity along the intracellular, P2Y12R-labeling along the extracellular line). After normalizing and scaling, microglial contacts were identified along the membrane of the endothelial cell, where microglial fluorescent intensity was over 20% of the maximal value, for at least along a 500 nm long continuous segment. Then the contact area was extended with 500-500 nm on both sides, and TOM20 fluorescent intensity within these areas was measured for “contact” value. TOM20 fluorescent intensity outside these identified contacts was considered “non-contact”. For the analysis of GFAP^+^ astroglial cell body contact frequency, CLSM stacks with double immunofluorescent labeling (GFAP and P2Y12R) were acquired from mouse cerebral cortex. All labeled and identified astrocytes were counted when the whole cell body was located within the Z-stack. To assess microglia process motility, baseline (28 min) and after 3x CCAo (49 min) two-photon image sequences were exported from MES software v.5.3560 (Femtonics Ltd., Budapest, Hungary) and analysed using FIJI (version 2.0.0, NIH, USA). The acquired hyperstacks were motion corrected using the StackReg plugin, then individual perivascular microglia processes (30 processes/image/plane) were tracked using the Manual Tracking plugin of FIJI. Based on the obtained XYZ coordinates, process motility speed was calculated. To study the effects of 3xCCAo on microglial morphology, 3-3 C57BL/6J mice were randomized into two groups: CCAo, or sham surgery. 24 hours after 3x CCAo, mice were transcardially perfused and processed for automated microglial morphology analysis. In both cases, 100μm thick sections with microglia (Iba1) and cell nuclei (DAPI) labeling were imaged with CLSM (0.2μm/pixel, Z-step of 0.4μm). Obtained confocal stacks were processed with the Microglia Morphology Quantification Tool^91^. For LSCI recordings, venous sinuses were excluded from the analysis. LSCI generates relative perfusion values (arbitrary units) therefore CBF was expressed as percentage change over baseline values in the 3x CCAo experiments. For the CCA occlusion experiments, a 1 min long period (typically 250 datapoints), recorded at the beginning of the imaging session, was averaged and considered as baseline. Then, every occlusion and reperfusion event was normalized to baseline. To assess the plasticity of the cerebrovasculature in response to repeated hypoperfusion, normalized occlusion or reperfusion events were averaged and compared between experimental groups. To demonstrate the CBF kinetics of individual animals, every 20th image was extracted and CBF values presented on a scatter plot (Supplementary Fig.4b). Quantification of P2Y12R immunostaining in control and microglia depleted tissues or P2Y12R and CD206 immunostaining in control and clodronate-treated mice were performed in at least three, randomly selected fields of view within the cortex, on three different coronal planes in each mouse. Data obtained from every mouse brains were averaged and compared between experimental groups. To investigate microglial actions on hypercapnic vasodilation, two photon image sequences were exported from the MESc software v.3.5.6.9395 (Femtonics Ltd.). After motion correction using the StackReg plugin of FIJI, the extent of vasodilation was measured and expressed as percentage of baseline at maximal vasodilation using FIJI. Obtained data were averaged and compared between control and microglia depleted or between CX3CR1^GFP/+^ and CX3CR1^GFP/+^ × P2Y12R KO mice. The hypercapnia-evoked CBF responses (2 min long) were normalized to the baseline (1 min long), then maximum values of individual responses were averaged per animal and compared between control and microglia depleted groups. Concentrations of released cellular purine nucleotides in response to hypoxia or hypercapnia were calculated by a two-point calibration curve, using internal standard method. The obtained values were averaged and compared to baseline ones. For neurovascular coupling experiments (manual and electromechanical whisker stimulations) the stimulus-evoked responses were normalized to baseline and were expressed as CBF increase (% change). The magnitude of evoked CBF responses was compared between experimental groups. GCaMP6s signals of individual neurons were collected with MESc software v.3.5.6.9395 (Femtonics Ltd.) and imported into the MES software v.5.3560 (Femtonics Ltd.) curve analysis module. The individual cellular [Ca^2+^]^i^ traces were normalized to the baseline GCaMP6s signal and data were derivated to relative fluorescence intensity change (ΔF/F). Then area under the curve (AUC) was calculated for each response, and AUC values were compared between experimental groups. During electrophysiological assessment, the baseline frequency of individual units was determined by averaging a 5min long period at the beginning of registration, when whisker stimulation was not applied. Only those units were selected for further analysis, which responded to electromechanical stimulations. Then the stimulus-evoked responses were corrected to baseline frequencies (called as baseline corrected response frequency) and the magnitude of responses was compared between experimental groups.

## Statistical Analysis

Animals were randomized for *in vivo* experiments using GraphPad Random Number Generator. Sample size was determined by a priori power calculation using G*Power 3.1.9.2 with mean differences and 20-25% standard deviations based on pilot studies (power 80%, α=0.05). Data were analyzed by GraphPad Prism 7.0 software, unless stated otherwise. Data were assessed for normal distribution using the D’Agostino-Pearson normality test or the Shapiro-Wilk W-test in order to determine parametric or non-parametric analysis. For comparing two or more groups with normal distribution the unpaired t-test with Welch’s correction either one-way ANOVA with Dunett’s multiple comparison test or two-way ANOVA with Tukey’s or Sidak’s multiple comparison test was used. For unevenly distributed data, the Mann-Whitney test either one-way ANOVA with Dunett’s multiple comparison test or two-way ANOVA with Tukey’s or Sidak’s multiple comparison test was used. Please, refer to the figure legends and the results section concerning the actual study design.

## Acknowledgements

The authors would like to thank László Barna, Csaba Pongor and Pál Vági the Nikon Microscopy Center at the Institute of Experimental Medicine, Hungary and Auro-Science Consulting, Ltd., for kindly providing microscopy support. We also thank the Medical Gene Technology Unit and the Cell Biology Center at the Institute of Experimental Medicine for their support. Functional ultrasound imaging was carried out at the ElfUS core facility, part of the IPNP, Inserm 1266 unit and Université de Paris. The authors are grateful to Plexxikon for providing PLX5622 for these studies. The authors would also like to thank the Department of Pathology, St. Borbála Hospital, Tatabánya, and the Human Brain Research Lab at the Institute of Experimental Medicine (Zsófia Maglóczky) for providing human brain tissue. The authors would like to acknowledge Dóra Gali-Györkei for her excellent technical assistance.

## Funding

This work was supported with funding from the ERC-CoG 724994 (AD), ERC-StG 715043 (BH), the ‘Momentum’ Program of the Hungarian Academy of Sciences (LP2016-4/2016 to A.D and BH), “PurinesDx” (BS), the Hungarian Brain Research Program [KTIA_13_NAP-A-I/2 (AD) and CCNKFIH KH125294 (BH). This project has also received funding from the European Union’s Horizon 2020 research and innovation programme under the Marie Skłodowska-Curie grant “ENTRAIN” agreement No. 813294 (ARB and AD). K.SZ and D.M. received support from the European Union’s Seventh Framework Programme (FP7/2007–2013) under Grant Agreements HEALTH-F2-2011-278850 (INMiND), FP7 HEALTH-305311 (INSERT), the Thematic Excellence Programme (TKP) of the Ministry of Innovation and Technology of Hungary, the COST Action CA16122 and under Grant Agreement No 739593. NL., K.SZ and CC were supported by the János Bolyai Research Scholarship of the Hungarian Academy of Sciences. CC (UNKP-20-5) and BP (UNKP-20-3-II) were supported by the New National Excellence Program of the Ministry for Innovation and Technology. BH was a member of the FENS-Kavli Network.

## Author’s contributions

EC and NL designed and conducted the experiments; EC, NL, CC, ZK, RF, BP, ADS, JCM, AK, LH, MB, ÁM performed imaging, light microscopy, histology and data analysis; CC designed anatomical experiments and performed electron microscopy; DS and KS contributed to SPECT and PET imaging; NL, ZK and ARB conducted cell culture studies; ÁM and EF designed and performed the CCAo study with brain pH measurements; ZsL, AK and JCM designed and performed functional ultrasound measurements; DB and BH performed electrophysiology, data analysis and provided intellectual support; BLW ZsL, ZB and EF contributed reagents and provided intellectual support; MB and BS performed HPLC analyses and provided intellectual support; AD devised the study, designed the experiments, provided overall supervision for the project, obtained funding and wrote the manuscript with input from all authors.

## Competing interests

B.L.W. is a past employee of Plexxikon.

## Data and materials availability

All data needed to evaluate the conclusions in the paper are present in the paper and/or the Supplementary Materials. Additional data related to this paper may be requested from the authors.

## Notes

### Competing Interest Statement

The authors have declared no competing interest.

