## Supplementary Material for "Microglia control cerebral blood flow and neurovascular coupling via P2Y12R-mediated actions"

| Subject | Code | Gender | Age (years) | Health status | Comorbidities | Cause of death | Tissue sample type |
|---|---|---|---|---|---|---|---|
| Control subject | SKO13 | female | 60 | normal | chronic bronchitis | respiratory arrest | free floating and paraffin sections |
| Control subject | SKO16 | male | 73 | normal | Unspec. atherosclerosis, pneumonia | respiratory arrest | free floating and paraffin sections |
| Control subject | SKO20 | male | 27 | normal | unspec. jaundice, malignant pancreatic neoplasm. | pulmonary embolism | free floating sections |

**Supplementary Table 1.** Patient data and processing of post-mortem human brain tissues.

### Supplementary Figures

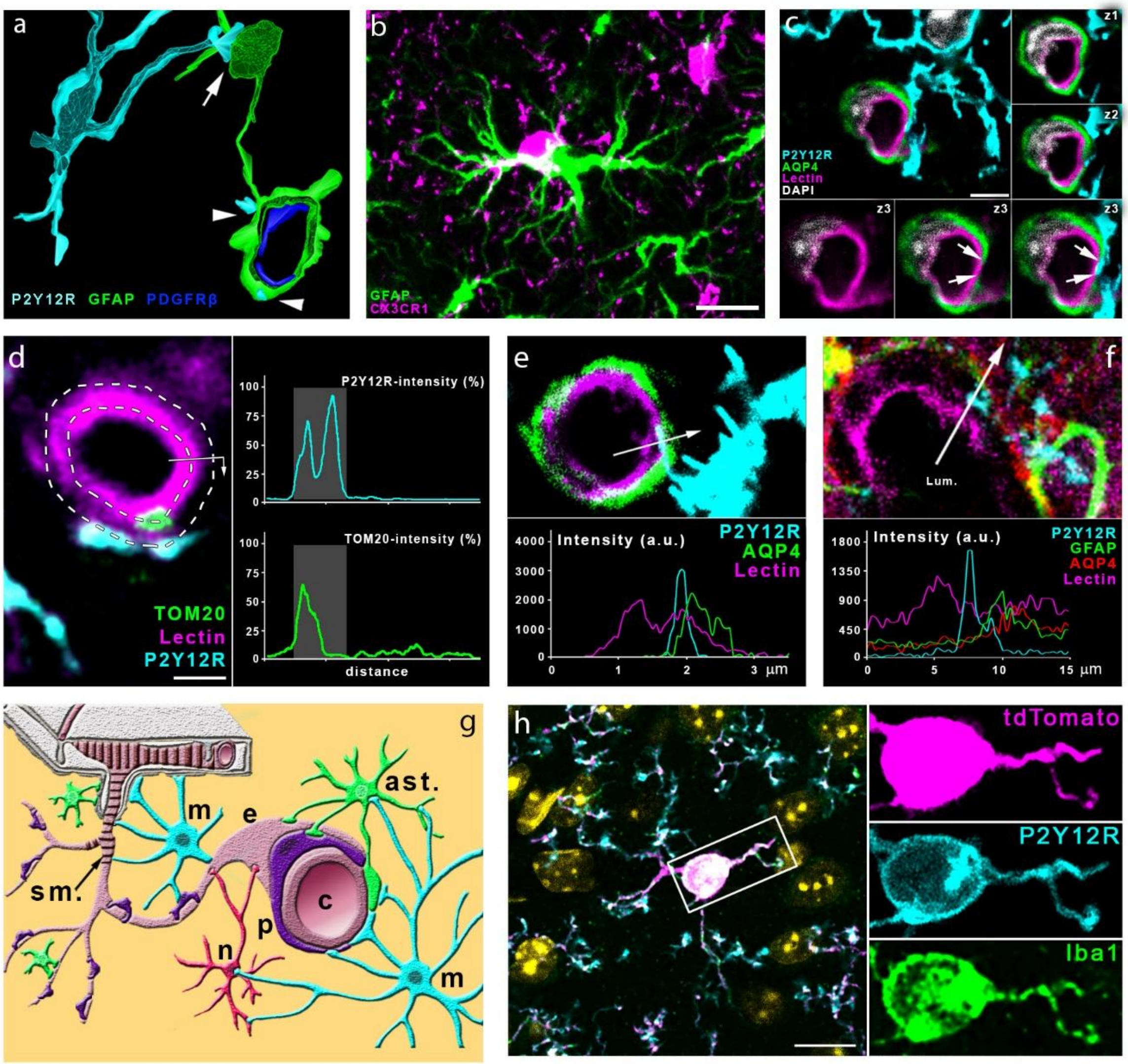


**Supplementary Figure 1. a)** 3D reconstruction of a high resolution CLSM Z-stack shows microglia (P2Y12R, cyan) contacting both the cell body of an astrocyte (GFAP labeling, green, arrow) and astrocytic endfeet (arrowheads) ensheathing a capillary. Pericytes are visualized by anti-PDGFRβ labeling (blue). **b)** Microglia (CX3CR1, magenta) are capable to form direct contact by their cell body with astrocytes (GFAP labeling, green) in the cerebral cortex. Scale bar: 10 µm. **c)** CLSM image show P2Y12R-positive microglial process (cyan) contacting perivascular aquaporin-4 (AQP4)-positive astrocyte endfeet (green) and also extends to the endothelial layer (Lectin, magenta) where astrocytic coverage is not present (arrows). z1-z3 panels show the contact area on three consecutive confocal sections. Scale bar: 3 µm. **d)** The process of semi-automated unbiased analysis of fluorescent intensity area for the graph presented in Fig.1J is depicted. White dashed lines represent the outer and the inner profiles, based on the outline of the endothelial cell. P2Y12R intensity was measured along the outer, TOM20 intensity along the inner profile, starting from the arrow. The intensity values are plotted (right) along the perimeter of the vessel. Contact site (marked by the grey column in the plots) was defined automatically. Scale bar: 2 µm **e)** CLSM image and fluorescent intensity plots show microglial process extending beyond perivascular astrocytic endfeet to interact with the endothelium. The fluorescent intensity profile plot (measured along the 3.5 um long white arrow) clearly shows the presence of the microglial process under astrocyte endfeet. **f)** CLSM image and fluorescent intensity plots show microglial processes interacting with GFAP- and AQP4-positive astrocytes in the human brain. The fluorescent intensity profile plot (measured along the 15 µm long white arrow) clearly shows the presence of the microglial process between the endothelium and the endfeet of perivascular astrocytes. **g)** Schematic summary of contacts formed between microglia (cyan, m) and different cell types of the neurovascular unit. Neurons (red, n), astrocytes (green, ast.), pericytes (purple, p), endothelial cells (pale crimson, e) and vascular smooth muscle cells (dark crimson, s.m.) are shown. **h**) Characterisation of CX3CR1$^{tdTomato}$ mice. Parenchymal tdTomato-positive cells co-express Iba1 and P2Y12R in the cerebral cortex. Cell nuclei stained with DAPI appear in yellow pseudocolor in the merged image. Scale bar: 10 µm.

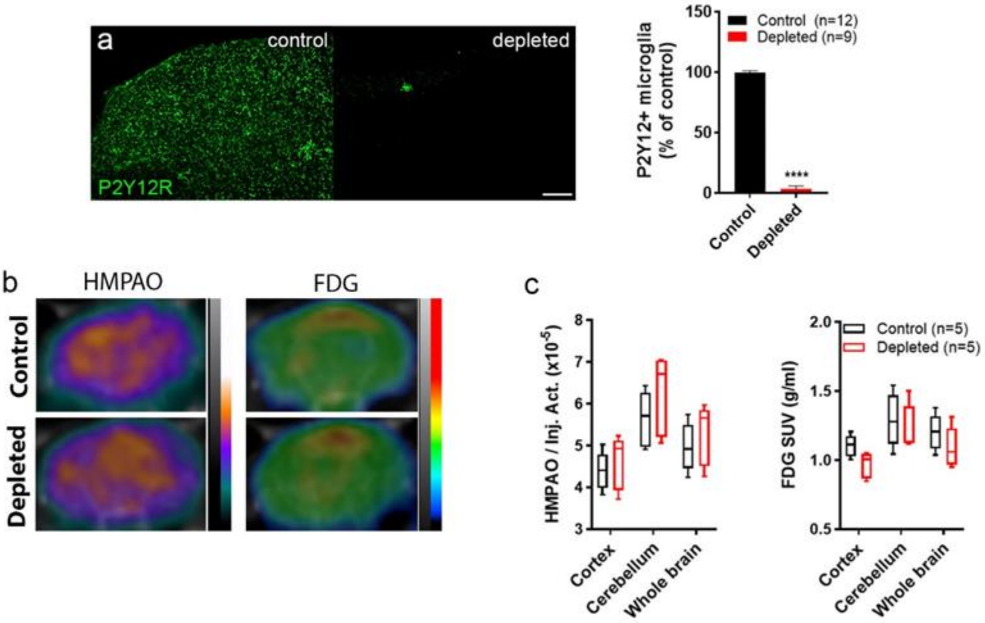


**Supplementary Figure 2. a)** Feeding C57BL/6J mice with a chow diet containing PLX5622 results in an almost complete (97%) elimination of resident microglia as evidenced by the numbers of P2Y12R-positive cells in the cerebral cortex. **b-c)** [99mTc]-HMPAO SPECT and [18F]-FDG PET images of control and microglia depleted mice. Proportion of measured and injected HMPAO activity and standard uptake values (SUV) of FDG are shown. Atlas-based region of interest analysis (b) shows no significant differences between the normalized regional uptake values (c) of the two groups. Data are expressed as mean ± SEM. n=12 control and n=9 depleted mice per group, ****$p<0.0001$ control versus depleted, unpaired t-test with Welch's correction (a); n=5- 5 mice, *p control vs depleted, two-way ANOVA followed by Sidak's multiple comparison test (c). Scale bar: 100 µm.

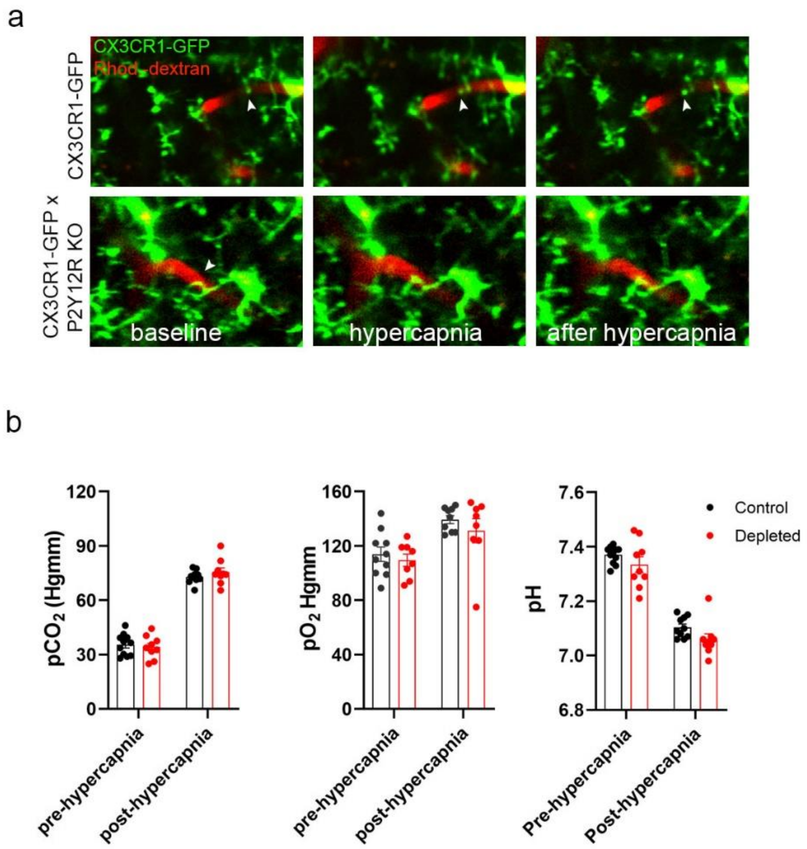


**Supplementary Figure 3. a)** *In vivo* two-photon imaging was performed with resonant scanning (32Hz) in the somatosensory cortex of CX3CR1$^{GFP/+}$ x P2Y12R KO and CX3CR1$^{GFP/+}$ (P2Y12R-competent) mice following intravenous Rhodamine B-Dextran (Rhod.-dextran) administration to visualize blood vessels. After recording 60 seconds of baseline, vasodilation was induced by inhalation of 10% $CO_2$ in air for 120 seconds (61-180 s) under normoxic conditions, followed by 60 seconds of post-hypercapnia recording, using identical protocol to that shown in Fig.4a-d,i. **b)** Arterial $pCO_2$, $pO_2$ and pH measurements under ketamine-medetomidine anesthesia after the administration of atipamezole, performed before and after hypercapnic challenge. Blood samples were taken from the femoral artery. No significant difference was observed between control and microglia-depleted mice. Data are shown as mean ± SEM. Two-way ANOVA followed by Sidak's multiple comparison test (b), n=10 control and n=8 depleted mice (b).

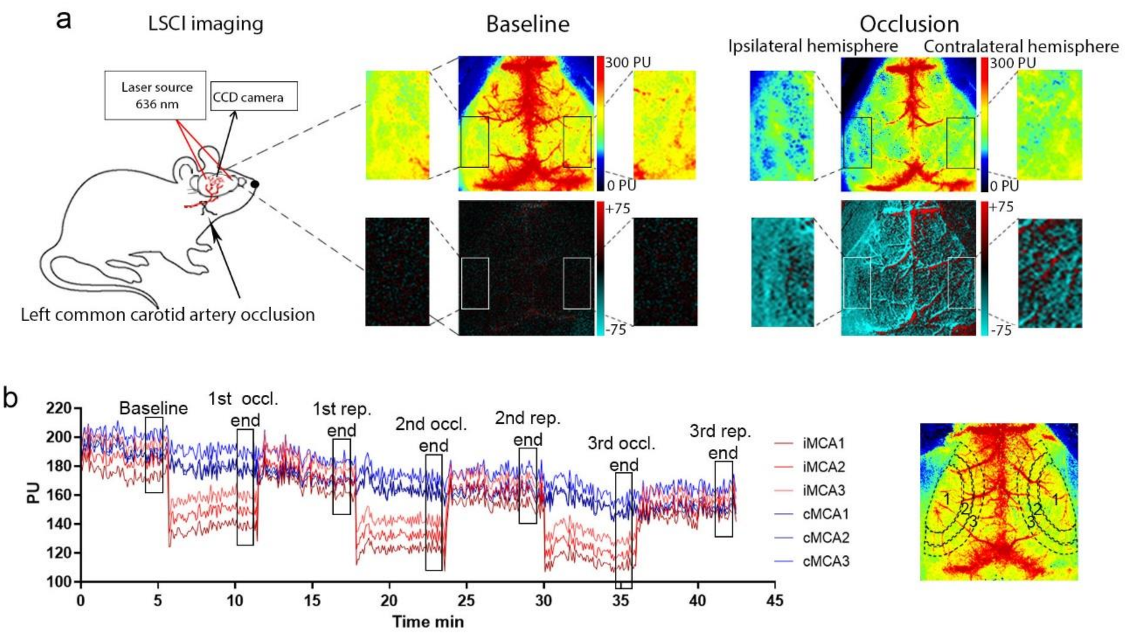


**Supplementary Figure 4. a)** CBF was measured during transient left CCA occlusion through the intact skull bone by LSCI. Representative perfusion (0 - 300 PU, on the top of panel a) and difference images (-75 - +75, on the bottom of panel a) show baseline CBF and perfusion changes during CCA occlusion. **b)** Representative graph showing the typical kinetic of repeated (3x CCA) occlusions on the areas (MCA1-3 areas) investigated on both hemispheres. ROIs are shown on a representative perfusion image on the right. Black rectangles on the kinetic graph display the sections of curves, which were used for detailed analysis.

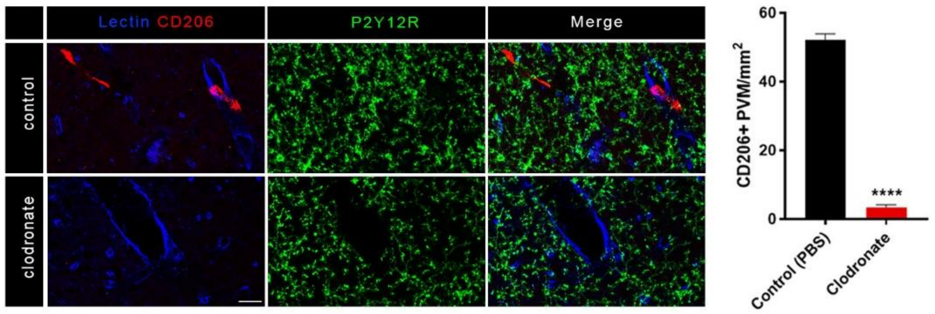


**Supplementary Figure 5.** Intracerebroventricular (ICV) clodronate administration resulted in the depletion of CD206-positive perivascular macrophages (PVMs), but did not affect microglial cells (P2Y12R labeling, green). Blood vessels were visualized using the endothelial marker, Tomato lectin (blue), scale bar: 20 µm. Quantification of the number of PVMs after ICV clodronate liposomes or PBS injection. Unpaired t-test with Welch's correction, n=5-5 mice per group, ****p<0.0001 clodronate versus control.

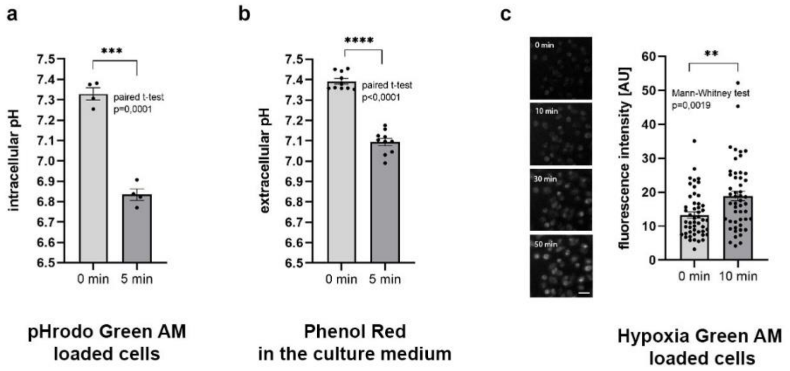


**Supplementary Figure 6**. **Validation of *in vitro* hypoxia and hypercapnia. a-b)** Both intracellular (a) and extracellular (b) pH markedly decreases within a few minutes after exposing cells to 15% $CO_2$/85% air gas mixture, as a model of hypercapnia. Extracell pH was determined by Phenol Red absorbance measurements while intracell pH was measured as changes in pHrodo Green AM dye fluorescence in glial cells. **c)** Hypoxia Green AM loaded cells exhibit significant increase in fluorescent intensity within 10 minutes after placing microglia cultures to hypoxic environment (1% $O_2$/5% $CO_2$/94% $N_2$). The reagent begins to fluoresce when oxygen level drops below 5%. Scale bar: 30 µm. Data are expressed as mean ± SEM; n=4 parallels per group, ***p=0.0001 0 min vs 5 min, paired t-test, (a); n=10 parallels per group, ****p<0.0001 0 min vs 5 min, paired t-test, (b); n=50 parallels per group, **p=0.0019 0 min vs 10 min, Mann-Whitney test, (c).

### Supplementary Video legends

**Supplementary Video 1.** Cerebral blood vessels were visualized by intravenous FITC-Dextran administration in CX3CR1$^{tdTomato}$ mice and microglial process dynamics was investigated by *in vivo* two-photon microscopy along the vascular tree. Please, refer to Figure 1.a-b for further details. Figure 1.b depicts identical blood vessels to those shown in the video file with arrowheads indicating the contact surfaces between microglial processes and 1$^{st}$ or 2$^{nd}$ order capillaries.

**Supplementary Video 2.** Representative Laser Speckle Contrast Imaging (LSCI) videos showing the whisker stimulation-evoked reduced neurovascular coupling response in microglia depleted mice compared to controls. Difference images are shown, which display the stimulus-evoked CBF increase over baseline. White rectangles indicate the area of the barrel cortex.

**Supplementary Video 3.** Representative resonant *in vivo* two-photon imaging videos showing individual neuronal $[Ca^{2+}]_i$ responses to electromechanically controlled whisker stimulation in control and microglia-depleted Thy1-GCaMP6s mice. Red dot indicates the onset of the individual stimuli (15s long) repeated twice with 40s intervals.

**Supplementary Video 4.** Representative *in vivo* two-photon imaging videos recorded by the resonant scanner showing reduced hypercapnia-induced vasodilation in P2Y12R KO (CX3CR1$^{GFP/+}$ x P2Y12R KO) mice compared to control (CX3CR1$^{GFP/+}$) mice. Blood vessels were visualized by administration of Rhodamine B-Dextran. Hypercapnia was induced at 60 s and maintained until 180 s with a 60 s long post-hypercapnic period. Identical fields of view are shown in Fig.4i.

**Supplementary Video 5.** Representative Laser Speckle Contrast Imaging (LSCI) video showing the third occlusion-reperfusion period in microglia depleted and control mice. Difference images display the marked CBF reduction caused by the 3$^{rd}$ occlusion over the 2$^{nd}$ reperfusion period. White ellipses show the area of the cerebral cortex (MCA territory) with significant CBF reduction after CCA occlusion. Note: to reduce the size of the video, a 2 minutes-long section from the 3rd occlusion and a 3 minutes-long section from the 2nd reperfusion have been removed. No significant CBF changes were seen during these periods.
